# Nucleolar endonuclease MBLAC2 orchestrates co-transcriptional ribosomal RNA processing and global transcriptional output

**DOI:** 10.64898/2026.07.31.742021

**Authors:** Luciano E Marasco, Rui Sousa-Luís, Anna-Sophia Maeckel, Rebecca Smith, Christopher J. Schofield, Nick J. Proudfoot

**Affiliations:** Sir William Dunn School of Pathology, University of Oxford, Oxford, UK; Gulbenkian Institute for Molecular Medicine, Universidade de Lisboa, Portugal; Cell Signalling and Cancer Laboratory St Vincent’s Institute of Medical Research, Fitzroy, Victoria 3065, Australia; Department of Chemistry and the Ineos Oxford Institute for Antimicrobial Research, 12, Mansfield Road, University of Oxford, Oxford, OX1 3TA, United Kingdom

## Abstract

The metallo-β-lactamase (MBL)-domain superfamily includes multiple nucleases that process distinct transcript classes. We show that human MBLAC2 acts as a ribosomal (r)RNA quality control factor, being nucleolar specific, chromatin-associated and required for co-transcriptional pre-rRNA processing. Notably MBLAC2 is functionally distinct from other MBL containing proteins, being required for maintenance of nucleolar architecture, cell cycle progression and proliferative fitness. Selective MBLAC2 depletion causes accumulation of pre-rRNA intermediates that lead to perturbed RNA polymerase (Pol) I-associated RNA homeostasis and nucleolar stress signalling. Nascent RNA profiling with long-read nanopore sequencing reveals extended and aberrant pre-rRNA species, consistent with defective processing and transcript release. Remarkably disruption of rRNA processing reduces global Pol II transcriptional output, even though the Pol II nuclear distribution remains unchanged across genomic compartments. Altogether, we define MBLAC2 as the nucleolar branch of an MBL-domain RNA surveillance network that couples rRNA quality control with transcriptional homeostasis.

## Introduction

The information flow from eukaryotic genes to functional RNA molecules relies on a panoply of diverse and intricately regulated RNA processing strategies. Among these, ribosomal RNA (rRNA) biogenesis stands out due to the exceptionally high demand for ribosomes required to sustain protein synthesis. This is especially true during cell cycle transitions when biosynthetic activity peaks. rRNAs represent by far the most abundant transcripts of the cell followed by replication-dependent histone mRNA and transfer RNA (tRNA), which are also produced in large quantities, but are transcribed by different RNA polymerases and processed through distinct molecular pathways. Whereas tRNA and 5S rRNA are synthesized by RNA polymerase III (Pol III), and histone mRNA by RNA polymerase II (Pol II), rRNA is transcribed by the dedicated RNA polymerase I (Pol I)^1,2^. This specialized polymerase is optimized for high-throughput processing from tandemly repeated rDNA arrays^3^ .This division of labour amongst RNA polymerases underscores the evolutionary pressure to maintain dedicated and high-fidelity transcription systems for different RNAs whose abundance and timing are tightly coupled to cellular proliferation^4,5^.

Ribosomal RNA plays fundamental structural and functional roles in the ribosome. It acts as a scaffold for the assembly of ribosomal subunits and forms key functional centres in both the decoding centre of the small subunit (SSU) and the peptidyl transferase centre of the large subunit (LSU). The latter catalyses peptide bond formation. In the higher eukaryotic genome, rRNA genes are organized into tandem arrays, each unit comprising the 18S, 5.8S, and 28S rRNA coding regions separated by internal and external transcribed spacers (ITS and ETS). These spacers are present in the 47S rRNA precursor and must be removed through tightly coordinated RNA processing events, involving both endonucleolytic and exonucleolytic cleavage. While many of the factors involved in this complex processing cascade have been elucidated in yeast and other model eukaryotes, the identity and regulation of analogous enzymes in vertebrate cells remains incomplete^6,7,8^.

A central feature of RNA metabolism is that each transcript class is coupled to dedicated processing and quality-control mechanisms. Several of these pathways are executed by proteins containing metallo-β-lactamase (MBL) fold domains, a structurally conserved family of metal-dependent enzymes that includes multiple DNA and RNA nucleases, surveillance factors, and detoxification enzymes such as hydroxyacylglutathione hydrolase (HAGH) and the persulfide dioxygenase ETHE1^9,10^ (Fig. 1A). Within this family, CPSF73 catalyses cleavage of pre-mRNA and replication-dependent histone transcripts^11,12^, INTS11 promotes processing and attenuation of noncoding RNA^13–16^, ELAC2 functions in mitochondrial tRNA maturation^17^ and MBLAC1 has been linked to histone mRNA metabolism^18^. This suggests that MBL-domain proteins have evolved as transcript class-specific RNA processing factors. However, whether there is an MBL nuclease dedicated to human pre-rRNA quality control has until now remained unclear.

**Figure 1.**
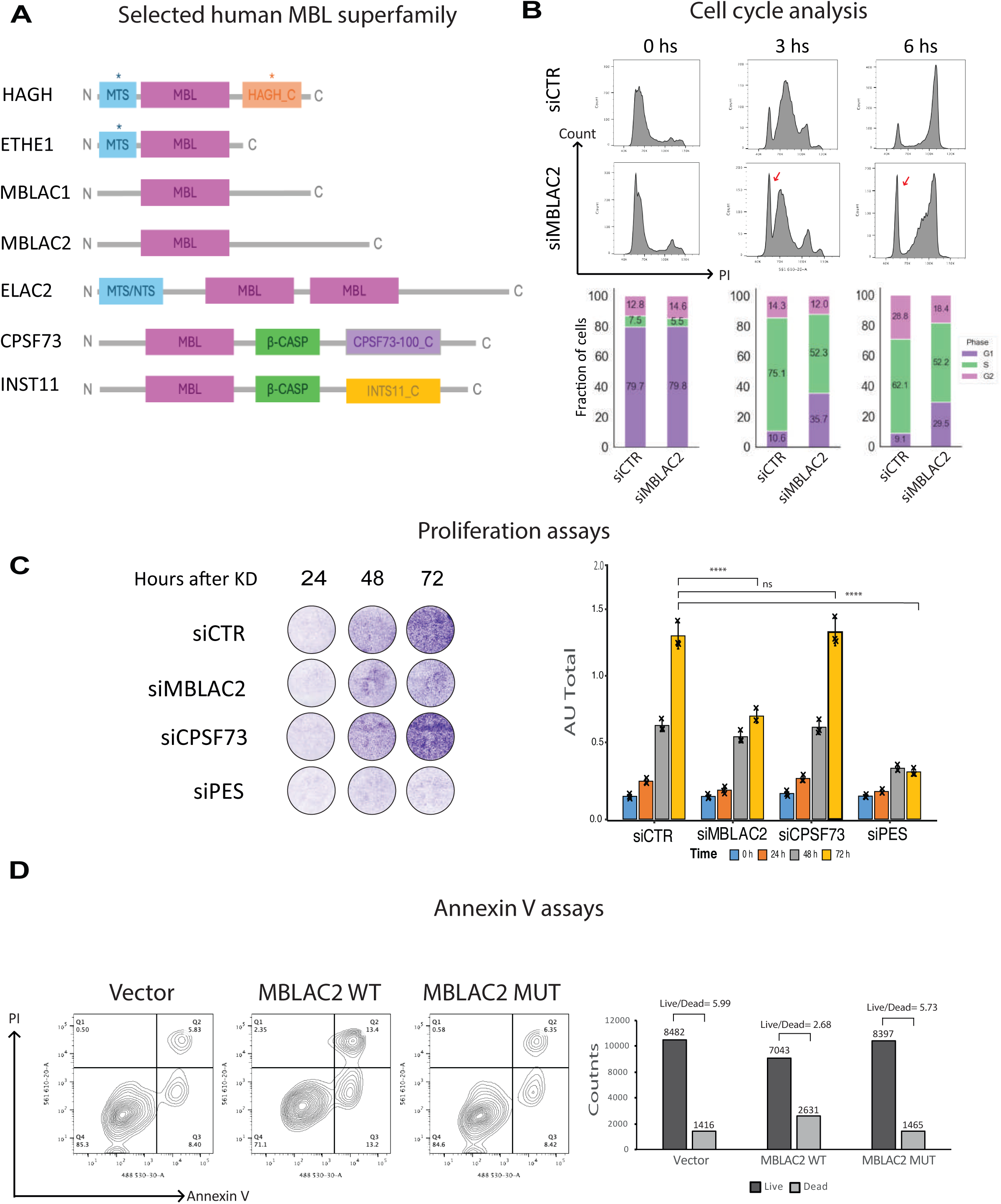
MBLAC2 is a member of the metallo-β-lactamase family and supports cell- cycle progression and cell viability. (A) Schematic representation of selected human metallo-β-lactamase (MBL)-domain proteins. HAGH, ETHE1, MBLAC1, MBLAC2, ELAC2, CPSF73 and INTS11 are shown with their annotated or predicted functional domains. MTS, mitochondrial targeting sequence; NTS, nuclear targeting sequence; β-CASP, β-CASP domain. (B) Cell-cycle progression following MBLAC2 depletion. HCT116 cells transfected with control siRNA or siMBLAC2 were synchronized by double-thymidine block and released for 0, 3 and 6 h. DNA content was measured by propidium iodide staining and flow cytometry. Red arrows indicate the G1-phase cell population. Representative histograms are shown above. Stacked bars show the proportion of cells in G1, S and G2/M. (C) Crystal-violet proliferation assay following depletion of MBLAC2, CPSF73 or PES1. Cells were collected 24, 48 and 72 h after siRNA transfection. Representative wells are shown at left and quantification of solubilized crystal-violet signal is shown at right. PES1 depletion was included as a positive control for impaired ribosome biogenesis. (D) Annexin V/propidium iodide flow-cytometry analysis of cells expressing empty vector, wild-type MBLAC2, or the catalytically impaired MBLAC2 mutant. Representative contour plots are shown on the left. Quantification of live cells (Q4) and dead cells, defined as the combined necrotic (Q2) and apoptotic (Q3) populations, is shown on the right; Q1 events were excluded. All experiments were performed in three independent biological replicates and data are presented as mean ± s.e.m. Cell-cycle data were analysed by two-way ANOVA with Šídák’s multiple-comparisons test, proliferation data using a two-factor linear model with HC3 robust standard errors and Holm–Bonferroni correction, and Annexin V/PI data by one-way ANOVA with Tukey’s multiple-comparisons test. *P < 0.05; **P < 0.01; ***P < 0.001; ns, not significant.

rRNA synthesis and processing occur in the nucleolus, a large membrane-less nuclear organelle organized around active rDNA repeats^19,20^. Nucleolar morphology is highly sensitive to perturbations in ribosome biogenesis. Its disruption can arise not only from acute external stressors but also from intrinsic homeostatic imbalance. Classical stressors such as Actinomycin D, UV irradiation, heat shock or nutrient deprivation all induce nucleolar fragmentation and redistribution of nucleolar components throughout the nucleoplasm^20,21^. The accumulation of unprocessed pre-rRNA, resulting from inefficient cleavage, maturation or surveillance, can disrupt the molecular stoichiometry and spatial organization required for nucleolar liquid–liquid phase separation (LLPS), ultimately compromising nucleolar integrity^6,7,20^. These morphological changes are often reversible upon restoration of transcriptional and processing homeostasis, highlighting the dynamic nature of the nucleolus.

Emerging evidence indicates that the activities of RNA polymerases I, II and III are functionally coordinated with one another and with RNA-processing pathways, particularly during ribosome biogenesis^22,23^. In yeast and mammalian systems, rRNA processing is tightly coupled to Pol I elongation, ribosome assembly and transcriptional termination^3,24,25^. Accumulation of unprocessed transcripts can impair Pol I release or termination, resulting in readthrough transcription into intergenic spacer regions (IGS) and further inhibition of co-transcriptional processing^24,26^. This creates a feed-forward loop in which defects in rRNA processing exacerbate transcriptional dysregulation, compounding nucleolar stress^27^.

We show here that MBLAC2, a nucleolar MBL-domain protein is required for pre-rRNA processing and rRNA quality control in human cells. MBLAC2 is related by sequence to MBLAC1 and, like MBLAC1 lacks the bCASP domain present in many MBL domain nucleases, including CPSF73. ^9,10^ It is, however, functionally distinct from MBLAC1, localizes to nucleolar and chromatin-associated compartments, and is required to maintain nucleolar architecture, cell cycle progression and Pol I transcription regulation. Using nascent RNA profiling and long-read nanopore sequencing, we find that MBLAC2 depletion causes accumulation of unprocessed and extended pre-rRNA intermediates, supporting its role in co-transcriptional rRNA maturation. Moreover, perturbation of rRNA processing through loss of MBLAC2 alters global Pol II transcriptional output, without redistributing Pol II across genomic compartments. Taken together, our findings define MBLAC2 as the nucleolar branch of a broader MBL- domain RNA surveillance network. They reveal how rRNA processing status influences nucleolar integrity and transcriptional homeostasis in higher eukaryotes.

## Results

### MBL-domain proteins define transcript specific RNA processing pathways

Metallo-β-lactamase (MBL)-domain proteins constitute a functionally diverse superfamily of metal-dependent enzymes, some of which act as RNA nucleases in essential RNA processing and surveillance pathways^10^ (Fig. 1A). A striking feature of MBL family nucleases acting on RNA is that individual enzymes operate on distinct RNA classes within specific subcellular compartments. For instance, CPSF73 catalyses endonucleolytic cleavage during pre-mRNA 3′ end formation^12,28^. INTS11, a catalytic subunit of the Integrator complex, cleaves noncoding RNAs such as small nuclear RNAs (snRNAs), enhancer RNAs and promoter-proximal transcripts, thereby contributing to both RNA processing and transcriptional attenuation^13–15^. ELAC2 functions as a mitochondrial tRNA-processing ribonuclease^29^. MBLAC1 has been linked to replication-dependent histone RNA metabolism, suggesting that related MBL proteins may have evolved specialized RNA quality-control activities^18^. Despite this functional diversification, whether the MBL family contains a dedicated factor for human pre-rRNA processing and surveillance has until now remained unclear.

We tested MBLAC2 as a candidate MBL-domain protein that might act within the nucleolar branch of RNA quality control. This possibility was suggested by the requirement for endonucleolytic cleavage during maturation of the 47S pre-rRNA and by the still incomplete definition of vertebrate enzymes that process pre-rRNA intermediates^6,7^. To investigate whether MBLAC2 contributes to cellular homeostasis, we first assessed the consequences of its depletion on cell cycle progression. We synchronized HCT116 cells using a double thymidine block, depleted MBLAC2 by siRNA, and released them into the cell cycle prior to flow cytometric analysis (Supplementary Fig. 1A). Compared with control cells, MBLAC2-depleted cells accumulated in G1 and progressed more slowly through S phase at 3 and 6 h after release (Fig. 1B), indicating a defective G1/S transition. Consistent with this altered cell-cycle progression, MBLAC2 depletion was accompanied by deregulation of several G1/S-associated regulators such as Cyclin D and CDK4 and 6 (Supplementary Fig. 1B).

The cell cycle phenotype observed following MBLAC2 depletion was also observed for other MBL superfamily members. The MBL-domain protein HAGH, which is mainly located in the cytoplasmic, and ETHE1, which is in mitochondria, are unlikely to act in nuclear RNA processing, despite belonging to the MBL family. However similarly to MBLAC2 depletion, both HAGH and ETHE1 depletions altered cell-cycle profiles (Supplementary Fig. 1C). However, given their distinct subcellular localization and functions (Supplementary Fig. 1D), these effects are likely to reflect pathways different to that of MBLAC2. Because the available MBLAC2 antibodies did not provide reliable detection of endogenous protein, we quantified MBLAC2 mRNA by RT-qPCR. This confirmed efficient siRNA-mediated depletion at each time point examined after release. (Supplementary Fig. 1E). We also examined endogenous MBLAC2 expression across the cell cycle by re-analysis of a reported RNA-seq dataset^30^. MBLAC2 transcript abundance increased during progression through G1 and S phase and declined thereafter (Supplementary Fig. 1F). This temporal profile is compatible with the requirement for MBLAC2 during normal G1/S progression, although it does not in itself establish a direct role for MBLAC2 in cell-cycle regulation. Consistent with this cell cycle delay, MBLAC2 depletion also reduced cellular proliferation, as assessed by crystal violet assays (Fig. 1C). Notably, this phenotype was not observed upon depletion of CPSF73, whereas depletion of Pescadillo1 (PES1), a ribosome biogenesis factor required for large ribosomal subunit maturation, produced a stronger proliferative defect^31–33^. Thus, loss of MBLAC2 compromises proliferation in a manner more consistent with defective ribosome biogenesis than with general disruption of MBL- domain RNA nucleases.

We next tested whether the catalytic integrity of MBLAC2 influences cell fate. Overexpression of wild-type MBLAC2 rapidly increased Annexin V positivity, a clear marker for apoptosis. In contrast, a predicted catalytically impaired triple mutant, in which three conserved residues of the metallo-β-lactamase metal-binding pocket were replaced with alanine (H175A/D194A/H236A), produced a delayed and attenuated response (Fig. 1D). This observation suggests that the cellular consequences of MBLAC2 expression are linked, at least in part, to its nuclease activity. Because several MBL-family nucleases function as dimers or higher-order assemblies, the delayed phenotype of the mutant may reflect partial residual activity or interaction with endogenous MBLAC2. In contrast, depletion of MBLAC2 did not produce a marked apoptotic response under the conditions tested, suggesting that the proliferative defect observed after knockdown is not simply explained by acute cell death (Supplementary Fig. 1G). Consistent with this difference in Annexin V levels, wild-type MBLAC2 overexpression also reduced proliferative fitness in crystal violet assays, whereas the catalytically impaired mutant showed no comparable reduction (Supplementary Fig. 1H).

Together, these observations demonstrate that MBLAC2 depletion delays G1/S progression, reducing proliferation without causing acute apoptosis. Remarkably these cell cycle and proliferation effects are dependent on the integrity of the MBLAC2 MBL domain, suggesting an indirect RNA processing role.

### MBLAC2 is a nucleolar and chromatin-associated MBL protein

We show above that MBLAC2 depletion delays cell-cycle progression and reduces cell proliferation. We next examined whether this phenotype is associated with altered nucleolar organization. Immunofluorescence microscopy using UBF (Pol I upstream binding factor) and Nucleolin (NCL) as nucleolar markers showed that control cells contained multiple compact nucleolar structures per nucleus. In contrast, MBLAC2 depletion caused enlarged nucleoli and reduced the average number of nucleoli per nucleus (Fig. 2A–B). Notably, NCL staining frequently adopted a peripheral ring-like configuration surrounding a central NCL-poor region in MBLAC2-depleted cells. UBF staining also became more diffuse throughout the enlarged nucleoli, rather than remaining confined to discrete fibrillar centres, further indicating reorganisation of nucleolar sub-compartments. A similar pattern was observed following depletion of PES1 as a positive control, indicating a broader reorganisation of nucleolar sub-compartments under these conditions. Ring-like redistribution of nucleolar proteins has previously been associated with disruption of ribosome-biogenesis pathways and altered ribosomal processing^34^. Together, these data show that MBLAC2 depletion induces a marked reorganisation of the nucleolar architecture, affecting nucleolar number, size and internal sub-compartmental organisation.

**Figure 2.**
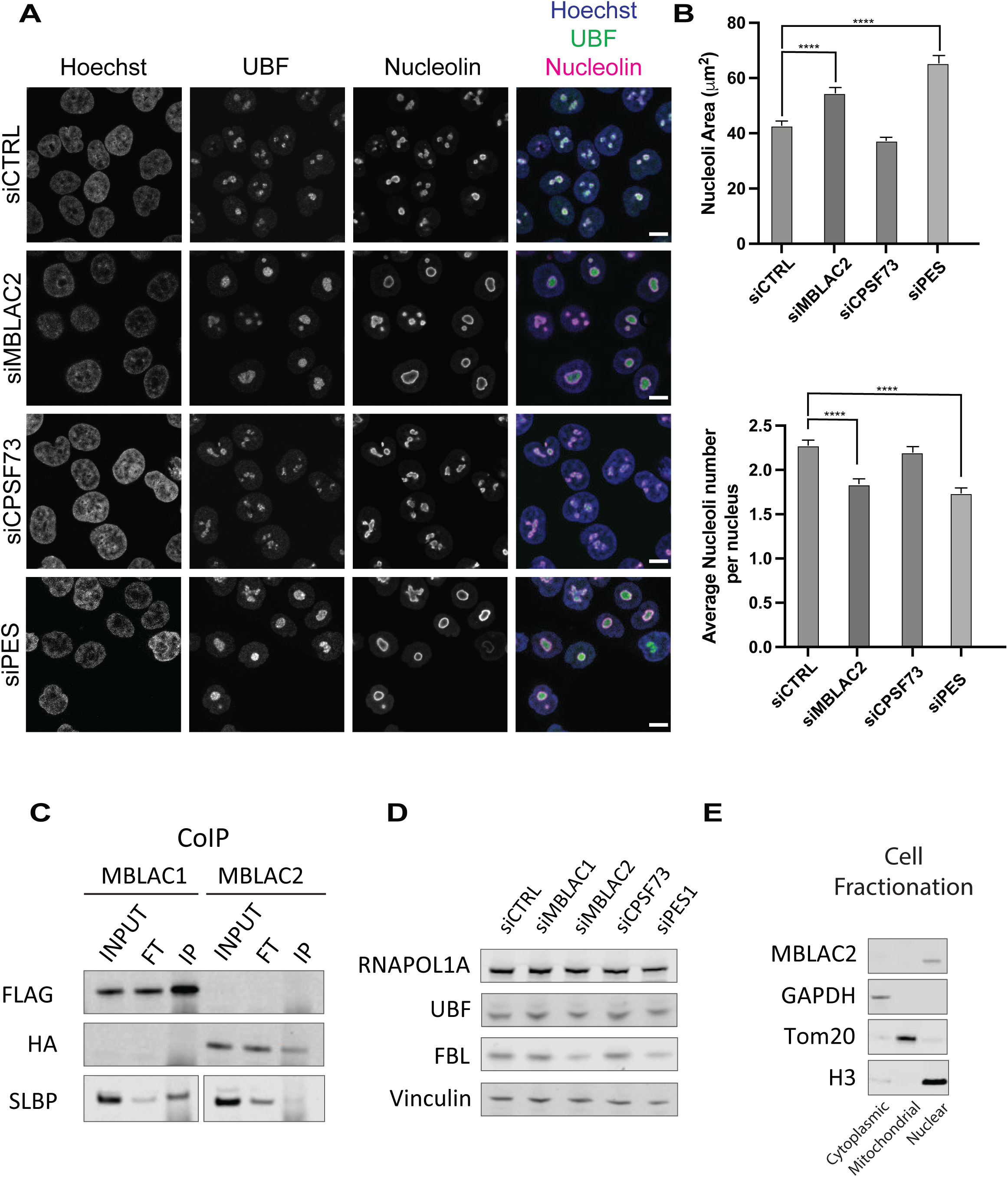
MBLAC2 is functionally distinct from MBLAC1 and is required for nucleolar integrity. (A) Immunofluorescence analysis of nucleolar morphology following depletion of MBLAC2, CPSF73 or PES1. HCT116 cells were stained for UBF and Nucleolin, and nuclei were counterstained with Hoechst. Representative images are shown. (B) Quantification of nucleolar area (top) and the average number of nucleoli per nucleus (bottom) from the experiment shown in (A). (C) Co-immunoprecipitation of epitope-tagged MBLAC1-FLAG or MBLAC2-HA followed by western blotting for FLAG, HA and SLBP. MBLAC1, but not MBLAC2, co- purified with SLBP under these conditions. Input, total cell extract; FT, flow-through; IP, immunoprecipitated fraction. (D) Western blot analysis of RNAPOL1A, UBF and fibrillarin (FBL) following depletion of MBLAC1, MBLAC2, CPSF73 or PES1. Vinculin was used as a loading control. (E) Subcellular fractionation of cells grown in a 150-mm culture dish into cytoplasmic, mitochondrial and nuclear fractions, followed by immunoblotting for MBLAC2. GAPDH, TOM20 and histone H3 were used as cytoplasmic, mitochondrial and nuclear fraction markers, respectively. For (A) and (B), data is a representative experiment of 3 independent biological replicates from between 209-385 cells per condition. Data were analysed by one-way ANOVA followed by Dunnett’s multiple-comparison test against siCTR. Bars show mean ± SEM. ****P < 0.0001; ns, not significant.

We next asked whether this nucleolar phenotype distinguishes MBLAC2 from its paralogue MBLAC1. MBLAC1 has previously been linked to replication-dependent histone RNA metabolism, through its association with stem-loop binding protein (SLBP), a factor required for histone mRNA 3′ end processing^35,18^. Native co-immunoprecipitation experiments with ectopic tagged proteins confirmed recovery of SLBP with MBLAC1, whereas MBLAC2 did not co-precipitate with SLBP (Fig. 2C). In accordance with this functional distinction, depletion of MBLAC1 had no effect on nucleolar area or nucleolar number, as judged by UBF and Nucleolin staining (Supplementary Fig. 2A-B). Thus, despite their related domain architecture, MBLAC1 and MBLAC2 appear to operate in distinct nuclear RNA-processing pathways.

Because MBLAC2 depletion altered nucleolar morphology, we next asked whether this phenotype reflected changes in the abundance of core nucleolar components. Western blot analysis revealed a modest reduction in fibrillarin (FBL) following MBLAC2 depletion, whereas the total levels of RNAPOL1A and UBF remained unchanged (Fig. 2D). MBLAC1 or CPSF73 depletion did not produce comparable changes. Together with the altered distribution of UBF and nucleolin observed by immunofluorescence, these results suggest that MBLAC2 depletion primarily causes a reorganisation of nucleolar architecture rather than a general loss of nucleoli or the Pol I transcription machinery. To further define the intracellular distribution of MBLAC2, we fractionated cells into cytoplasmic, mitochondrial and nuclear compartments. Endogenous MBLAC2 was detected predominantly in the nuclear fraction, consistent with its proposed role in nucleolar RNA metabolism (Fig. 2E). This nuclear enrichment is further supported by recent imaging showing that EGFP-tagged MBLAC2 localises constitutively to the nucleolus^36^.

Taken together, the data described above identify MBLAC2 as a nucleolar and chromatin-associated MBL protein that is functionally distinct from MBLAC1. This is shown by its lack of detectable association with SLBP, its enrichment in nuclear and chromatin fractions and the nucleolar defects caused by its depletion. These results are all consistent with a critical role for MBLAC2 in ribosome biogenesis rather than in histone RNA metabolism or in another MBL-domain nuclease pathway.

### MBLAC2 safeguards co-transcriptional pre-rRNA processing and rRNA homeostasis

The nucleolar role of MBLAC2 as shown by disruption of nucleolar architecture observed upon its depletion, suggests that MBLAC2 acts directly during ribosomal RNA biogenesis. Human rRNA is synthesized as a 13.3 kb or 47S precursor containing the mature 18S, 5.8S and 28S rRNAs separated by external (ETS) and internal transcribed spacers (ITS). Each rDNA repeat is further divided between the 47S transcription unit (TU) and a larger intergenic spacer (IGS) region. This latter region is partly transcribed by Pol II to generate long noncoding transcripts (lncRNA) as well as providing regulatory elements including Pol I promoter and termination sequences^22^. ETS and ITS RNA is removed by coordinated endonucleolytic and exonucleolytic processing^6,7^ so generating mature rRNA. Notably, transcription elongation by Pol I is tightly coupled to pre-rRNA processing and ribosome assembly, such that processing defects can feed back into polymerase dynamics^3,25,37^. To interrogate these steps, we measured rRNA transcript levels (both nascent and steady state) across the human rDNA repeat using the indicated 47S precursor RT/PCR primers including the 5′ETS, ITS1, ITS2, 3′ETS and intergenic spacer (IGS) regions, together with the positions of northern blot probes (Fig. 3A).

**Figure 3.**
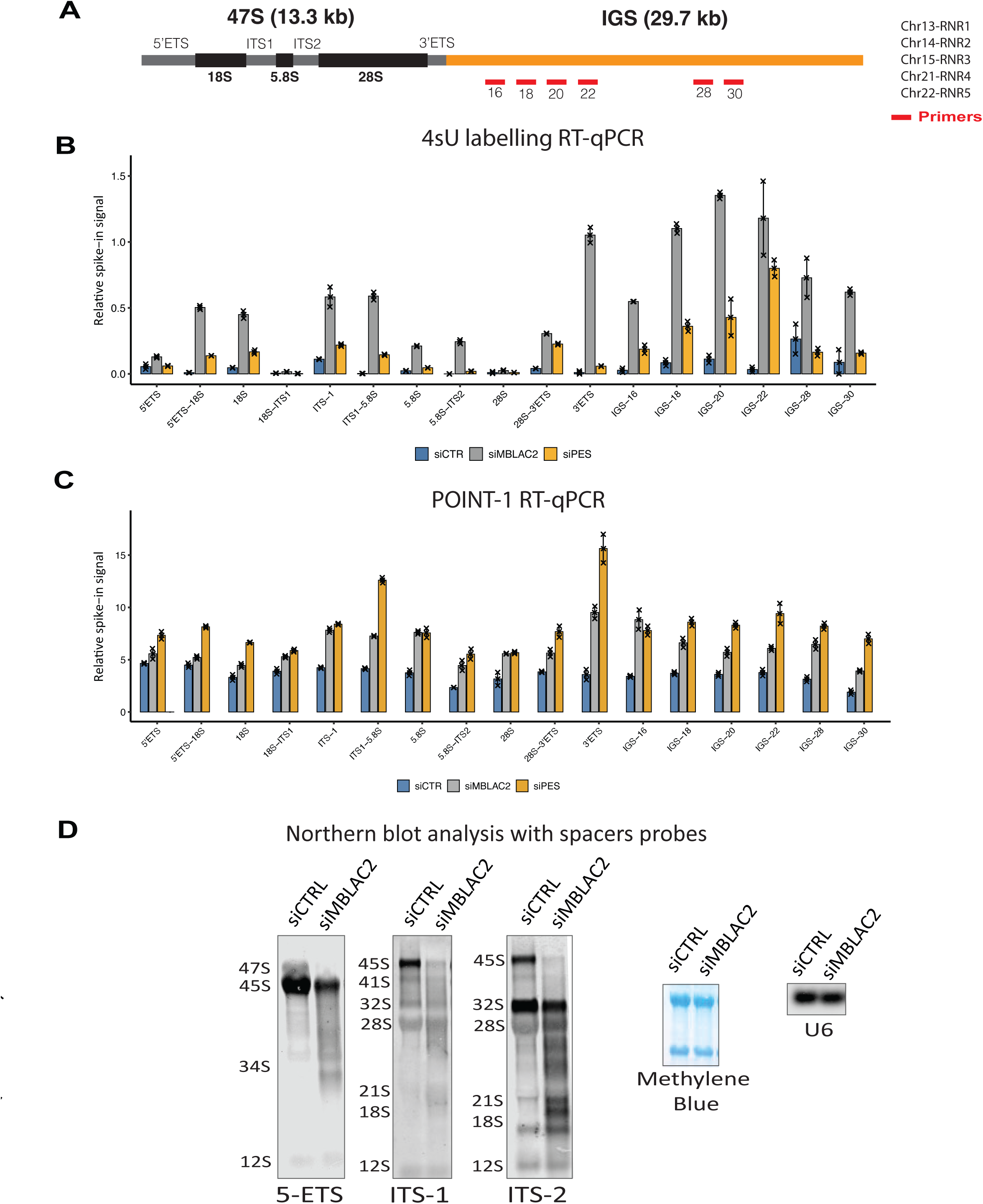
MBLAC2 depletion causes accumulation of nascent pre-rRNA intermediates and RNA polymerase I-associated readthrough transcripts. (A) Schematic representation of the human rDNA transcription unit and downstream intergenic spacer (IGS). The 47S pre-rRNA contains the mature 18S, 5.8S and 28S rRNA regions separated by the 5′ external transcribed spacer (5′ETS), internal transcribed spacers 1 and 2 (ITS1 and ITS2), and the 3′ETS. Amplicons used for RT-qPCR across the transcribed region and downstream IGS are indicated in red. (B) 4-Thiouridine (4sU) labelling followed by RT-qPCR across the rDNA transcription unit and downstream IGS in cells treated with siCTR, siMBLAC2 or siPES1. Nascent RNA levels were normalized to spike-in RNA. MBLAC2 depletion increased signal across spacer-containing pre-rRNA regions and downstream IGS positions. (C) POINT-1 RT-qPCR analysis of Pol I-associated RNA across the rDNA transcription unit and downstream IGS in cells treated with siCTR, siMBLAC2 or siPES1. Values were normalized to spike-in RNA. (D) Northern-blot analysis of pre-rRNA intermediates using biotinylated probes against the 5′ETS, ITS1 and ITS2 regions in control and MBLAC2-depleted cells. Positions of major pre-rRNA intermediates are indicated. Methylene-blue staining and U6 detection are shown as loading controls. For (B) and (C), data were analysed by two-way ANOVA with treatment and amplicon as factors, followed by Dunnett’s multiple-comparison test comparing each treatment with siCTR at the corresponding amplicon. Values represent mean ± SEM from n = 3 independent biological replicates. Northern blots are representative of n = 2 independent experiments. *P < 0.05; **P < 0.01; ***P < 0.001; ****P < 0.0001; ns, not significant.

We first tested whether MBLAC2 depletion affects newly synthesized rRNA precursors. For this, we performed 4-thiouridine (4sU) metabolic labelling followed by RT-qPCR across the rDNA locus. MBLAC2 knockdown caused accumulation of nascent spacer-containing pre-rRNA species, with increased signal across the 5′ETS, ITS1, ITS2 and 3′ETS regions (Fig. 3B). Nascent transcript levels were observed to substantially increase over downstream IGS regions, suggesting a major role for MBLAC2 in Pol I termination. PES1 depletion produced a related profile albeit with reduced effect. This is consistent with the established role of PES1 as a component of the PeBoW complex, which is required for pre-rRNA processing, maturation of the 28S rRNA, and large ribosomal subunit biogenesis^31–33^. Overall, these data indicate that depletion of MBLAC2 results in impaired maturation or turnover of newly synthesized pre-rRNA. It is remarkable that 4sU labelling is barely detectible in mature rRNAs. This underlines the fact that 4sU likely precludes correct RNA processing and folding of these abundant RNAs. Consequently, 4sU incorporation is mainly evident in unprocessed pre-rRNA.

We next asked whether defective processing is associated with altered Pol I associated RNA. To address this, we developed a new variation of the POINT-seq technique^38^ employing anti-Pol I rather than anti-Pol II antibodies called here POINT-1 (see Fig. 4A). This recovers RNA physically associated with Pol I and, as before, was measured by RT-qPCR using the same probes as above across the rDNA transcription unit. MBLAC2 depletion increased Pol I-associated RNA signal over the 47S precursor and within the IGS (Fig. 3C). In this case the effect was more pronounced after PES1 depletion. Notably, they both lead to accumulation of Pol I-associated pre-rRNA and downstream transcripts, consistent with inefficient processing, delayed transcript release or readthrough beyond the 3′ end of the rDNA transcription unit. This interpretation is in line with the view that rRNA processing, ribosome assembly and Pol I termination are all mechanistically coupled^37^.

**Figure 4.**
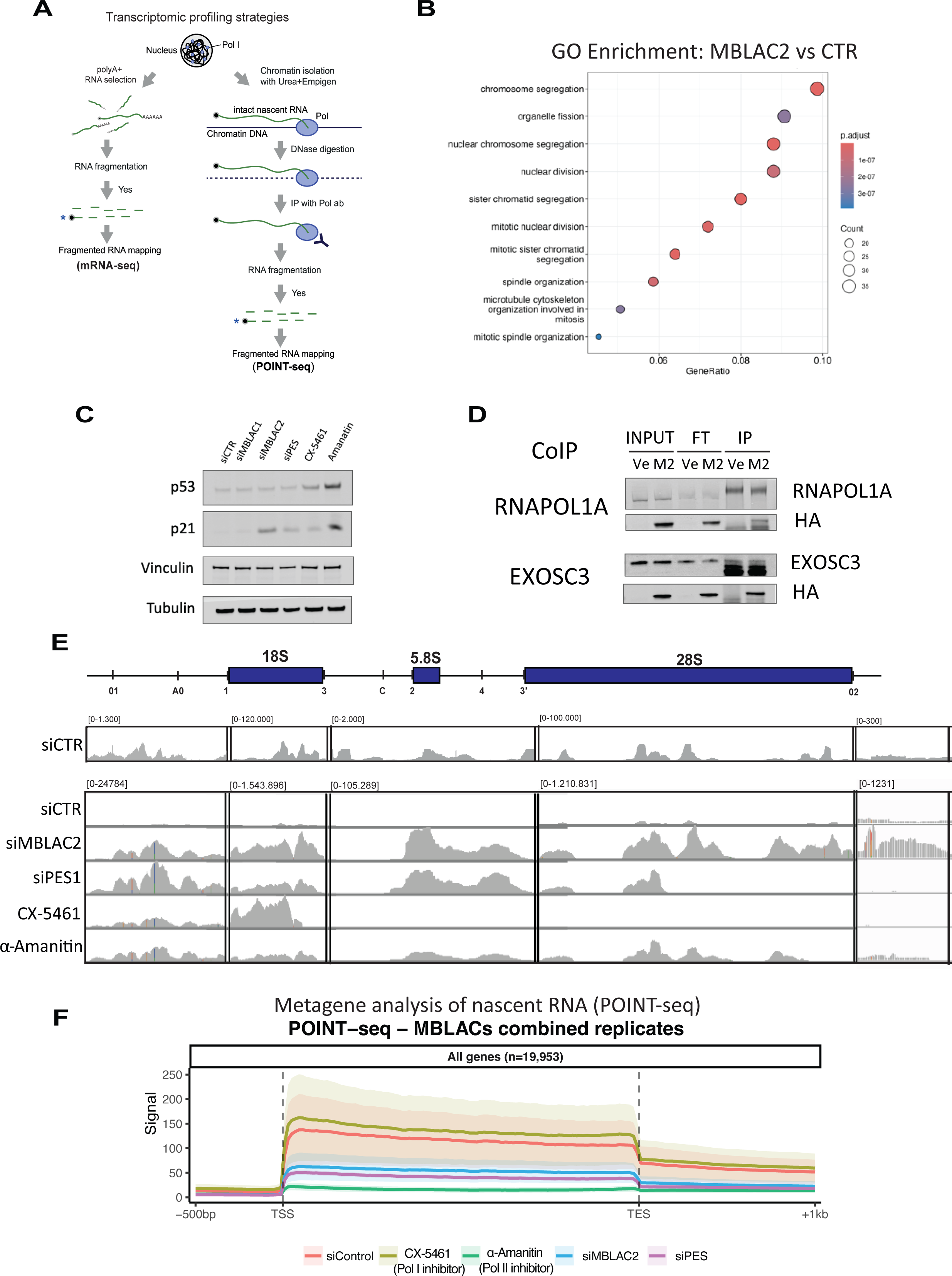
MBLAC2 depletion alters ribosome biogenesis pathways and reduces global Pol II transcription. (A) Overview of the transcriptomic profiling strategies used in this study. Poly(A) selected RNA sequencing was used to measure steady-state mRNA abundance, whereas POINT-seq was used to recover polymerase-associated nascent RNA from chromatin. The POINT-1 and POINT-2 datasets were subsequently used to examine Pol I-associated RNA across the rDNA repeat and Pol II-associated transcription across protein-coding genes, respectively. (B) Gene ontology enrichment analysis of differentially downregulated genes following MBLAC2 depletion relative to siCTR cells. Enriched terms include chromosome segregation, organelle fission, nuclear division and mitotic sister chromatid segregation. Dot size indicates the number of genes assigned to each term, and colour indicates the adjusted P value. (C) Western-blot analysis of p53 and p21 following depletion of MBLAC2 or PES1, or treatment with CX-5461 or α-amanitin. Vinculin and tubulin are shown as loading controls. (D) Co-immunoprecipitation of endogenous RNAPOL1A or EXOSC3 from cells expressing HA-tagged MBLAC2, followed by immunoblotting for the indicated proteins. HA–MBLAC2 was recovered in both RNAPOL1A and EXOSC3 immunoprecipitates. Ve, empty vector; M2, HA–MBLAC2; FT, flow-through; IP, immunoprecipitate. (E) Genome-browser profiles of POINT-1 signal across the human rDNA transcription unit following MBLAC2 or PES1 depletion or treatment with the Pol I inhibitor CX-5461 or the Pol II inhibitor α-amanitin. Mature rRNA regions and major processing sites are indicated above. The upper siCTR track is displayed using a library-specific scale, whereas the lower siCTR track is group-scaled together with the treatment tracks to allow direct comparison. (F) Metagene analysis of POINT-2 signal across annotated protein-coding genes, extending from 500 bp upstream of the transcription start site (TSS) to 1 kb downstream of the transcript end site (TES). Profiles represent the mean of two independent biological replicates per condition, and shaded areas indicate s.e.m. Differential-expression analysis of poly(A)-selected RNA-seq data was performed using DESeq2, with P values adjusted using the Benjamini–Hochberg procedure. Gene ontology enrichment was performed using clusterProfiler with false-discovery-rate correction.

Northern blots, using biotinylated probes as indicated, further support a role for MBLAC2 in pre-rRNA maturation. Thus, probes against spacer-containing regions reveal accumulation of pre-rRNA intermediates following MBLAC2 depletion, detected as “smeary” bands (Fig. 3D). These changes are consistent with inefficient processing of the transcribed spacers from the 47S precursor. In contrast, probes against the mature ribosomal RNA species gave less apparent unprocessed rRNA signal albeit aberrant isoforms were still observed (Supplementary Fig. 3A). Moreover, MBLAC2 depletion did not substantially affect global translation, as judged by puromycin incorporation in the SUnSET assay, whereas PES1 depletion reduced puromycin incorporation and served as a positive control for impaired ribosome biogenesis (Supplementary Fig. 3B). Transfected GFP expression was not substantially reduced after MBLAC2 depletion (Supplementary Fig. 3C). Given the long half-life of mature ribosomes, an immediate reduction in translation would not necessarily be expected following an early pre-rRNA processing defect and may only become apparent after prolonged MBLAC2 depletion. These observations all suggest that MBLAC2 acts primarily at early stages of pre-rRNA processing, before there is a noticeable decline in the production of mature or functional ribosomes. As a control for the specificity of the POINT-1 assay, we also tested whether immunoprecipitation of native Pol I occurs with co-purification of other nuclear RNA polymerases. Western blot analysis showed that Pol II and Pol III were present in the input and flow-through fractions but were not detectably recovered in the Pol I immunoprecipitate (Supplementary Fig. 3D). Thus, the RNA measured by POINT-1 analysis is unlikely to reflect contamination from Pol II or Pol III-associated transcripts and can be assigned more specifically to Pol I complexes.

Together, these data identify MBLAC2 as a nucleolar MBL-domain factor required for normal metabolism of nascent pre-rRNA. Its depletion causes accumulation of spacer-containing rRNA precursors and abnormal retention of Pol I-associated RNA into downstream rDNA regions, suggesting that MBLAC2 aids the coordination of pre-rRNA processing with Pol I transcript release as well as more generally in nucleolar homeostasis.

### rRNA processing status controls global Pol II transcriptional output without broad redistribution

Given that MBLAC2 depletion impairs pre-rRNA processing and disrupts nucleolar architecture, we next asked whether this nucleolar defect could extend to the broader transcriptional program of the cell. Ribosome biogenesis is tightly linked to cell growth^39^. Perturbations in rRNA synthesis or maturation can activate stress responses that extend beyond the nucleolus^20,21,39^. We therefore generated libraries from polyA-selected, mRNA-seq and POINT-isolated nascent RNA from Pol I (POINT-1-seq) and II (POINT-seq). This allowed us to compare steady-state gene expression with nascent transcriptional output at the genome-wide level.

mRNA-seq analysis showed that MBLAC2 depletion induces a transcriptional response compatible with impaired ribosome biogenesis and cell-cycle delay. Gene ontology analysis of differentially expressed genes revealed enrichment for cell-cycle-related terms, including chromosome segregation, organelle fission, nuclear division and mitotic sister chromatid segregation (Fig. 4B). Consistent with the G1/S phenotype described above, CDKN1A, encoding p21, was among the genes induced following MBLAC2 depletion (Table 1) and increased p21 protein abundance was also observed by western blotting (Fig. 4C)^40^. By contrast, a subset of genes involved in RNA metabolism and ribosome biogenesis was reduced (Table 1). Notably, EXOSC3, which encodes a core component of the nuclear RNA exosome, was among the downregulated transcripts. This observation suggests that loss of MBLAC2 may affect not only pre-rRNA processing but also RNA-surveillance pathways that normally act on aberrant or incompletely processed transcripts.

We therefore tested whether MBLAC2 associates with factors involved in Pol I transcription and RNA surveillance. Native co-immunoprecipitation experiments showed that ectopic expression of MBLAC2-HA recovered RNAPOL1A, the largest subunit of Pol I, and EXOSC3 (Fig. 4D). These interactions are consistent with MBLAC2 acting in close proximity to nascent Pol I transcripts and with a possible connection between pre-rRNA processing and exosome-linked RNA surveillance. PES1 depletion produced a related gene ontology profile, again dominated by cell-cycle and mitotic categories (Supplementary Fig. 4A). Although MBLAC2 and PES1 are distinct proteins, their depletion generates a similar cellular state associated with defective ribosome production and delayed cell-cycle progression.

We next examined Pol I-associated nascent RNA across the rDNA locus. POINT-1-seq profiles obtained by Illumina sequencing showed greatly increased signal following MBLAC2 depletion across the 47S transcription unit and into the downstream intergenic spacer regions as compared to the siRNA control (Fig. 4E). This suggests a high level of rRNA retention on the rDNA template especially of unprocessed rRNA transcripts. This is exacerbated following MBLAC2 depletion. siPES1 also gives a high signal over the 47S region, but not over the 3’ region half of the 28S region or into the IGS region. Apparently, there is a less severe 3’ end defect following loss of PES1 as compared to loss of MBLAC2. Overall, this confirms the proposal that depletion of either MBLAC2 or PES1 cause substantial retention of nascent rRNA across the 47S region. Notably, pharmacological inhibition of Pol I with CX-5461 eradicated the signal over the rDNA unit except for the promoter proximal region into the 18S gene. This residual promoter-proximal signal may reflect the TOP2B-inhibitory activity of CX- 5461, which could increase torsional stress ahead of Pol I and so stall elongation shortly after initiation^41^. The Pol II inhibitor α-amanitin which primarily inhibits Pol II, produced a diminished but similar profile to MBLAC2 depletion. This observation underlines the role of Pol II transcription in the control of Pol I gene expression levels as previously demonstrated^23^. In effect our POINT-1-seq results reinforce the view that upon MBLAC2 depletion, the relative contribution of reads mapping into the 3′ETS markedly increases. This redistribution is consistent with accumulation of Pol I-associated RNA near the 3′ end of the transcription unit and supports the view that defective pre-rRNA processing is associated with altered transcript release or termination. (Supplementary Fig. 4B).

Although rRNA is transcribed by Pol I, ribosome biogenesis also depends on Pol II and Pol III. Pol II produces mRNAs encoding ribosomal proteins and ribosome-assembly factors, as well as regulatory noncoding transcripts, whereas Pol III transcribes 5S rRNA and tRNAs, both required for ribosome function^1,2,5^. This functional interdependence implies that perturbation of pre-rRNA processing may be communicated to the other transcription systems. We therefore asked whether the MBLAC2-dependent nucleolar defect affects Pol II transcriptional output. The Pol II metagene profile was generated from POINT-2 libraries after normalization to total mapped reads/spike-in signal by averaging normalized signal across annotated protein-coding genes from TSS to TES. Remarkably, this analysis showed that MBLAC2 depletion drastically reduced global Pol II nascent RNA across protein coding gene bodies (Fig. 4F). A similar reduction was observed upon PES1 depletion. In contrast Pol I inhibition increased the Pol II metagene signal, implying a degree of molecular competition between Pol I and Pol II. As expected, Pol II inhibition by α-amanitin treatment removed most of the protein coding metagene signal. These data indicate that defective ribosome biogenesis severely restricts the overall output of Pol II transcription.

Importantly, the decrease in Pol II signal following MBLAC2 depletion was not accompanied by a major redistribution of Pol II reads across genomic compartments. In control cells, POINT-seq signal reads was predominantly intronic, with smaller reactions mapping to exons, TSS-proximal regions, TES regions, post-TES regions and intergenic regions. Upon MBLAC2 depletion, this global distribution was broadly preserved, with only modest changes in the relative contribution of each compartment (Supplementary Fig. 4C). Thus, MBLAC2 depletion appears to reduce the amplitude of Pol II transcription rather than extensively rewiring the genomic localization of Pol II.

This distinction is important. Acute Pol I inhibition lowers rRNA synthesis, whereas chronic defects in pre-rRNA processing or ribosome assembly, as produced by MBLAC2 or PES1 depletion cause accumulation of abnormal rRNA intermediates, nucleolar reorganization and transcriptional adaptation. Inhibition of Pol I, Pol II elongation by DRB or Pol II degradation by α-amanitin consistently produced similar effects on nucleolar structure as observed by nucleolar markers such as UBF and Nucleolin (Supplementary Fig. 4D). These observations suggest that Pol II transcription does not simply respond to the amount of rRNA being synthesized, but more directly to the functional state of nucleolar RNA metabolism.

Together, these results indicate that pre-rRNA quality control mediated by MBLAC2 is coupled to global transcriptional homeostasis. Loss of MBLAC2 triggers a ribosome-biogenesis stress program and reduces Pol II nascent transcription, while preserving the overall architecture of Pol II genomic distribution. Thus, MBLAC2 acts as a nucleolar MBL-domain factor required to maintain the processing competence of nascent pre-rRNA. When this function is impaired, the nucleolus enters a distinct stress state that is communicated to the Pol II machinery, revealing crosstalk between transcriptional machineries.

### Long-read sequencing reveals aberrant and extended pre-rRNA intermediates upon MBLAC2 depletion

The accumulation of spacer-containing pre-rRNA species detected by northern blotting, 4sU labelling and POINT-1 all suggest that MBLAC2 depletion alters the normal maturation of nascent Pol I transcripts. However, these approaches either interrogate selected positions across the rDNA repeat or rely on short-read coverage and therefore do not directly resolve the architecture of individual rRNA molecules. We therefore adapted POINT-1 for long-read nanopore sequencing of Pol I-associated nascent RNA, here termed POINT-nano (Fig. 5A). This approach preserves the continuity of individual RNA molecules and so allows pre-rRNA intermediates to be examined in the context of their start and end positions across the rDNA transcription unit^42^.

**Figure 5.**
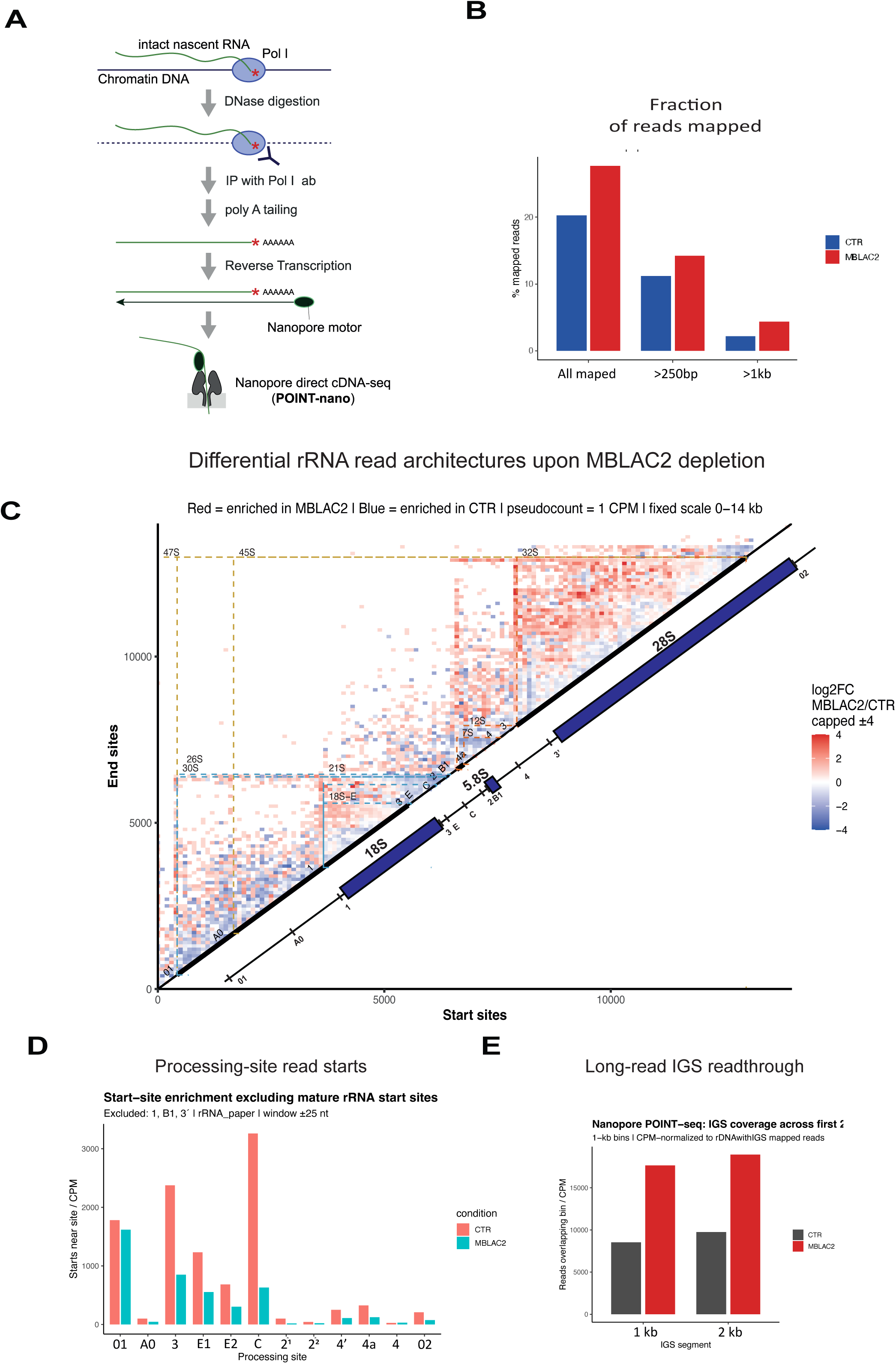
Long-read POINT-seq reveals aberrant pre-rRNA architectures and extended IGS-associated transcripts upon MBLAC2 depletion. (A) Schematic of the POINT-nano workflow. Pol I-associated nascent RNA was isolated from chromatin, polyadenylated, reverse-transcribed and analysed by nanopore direct cDNA sequencing. (B) Fraction of total input reads mapping to the rDNA reference in control and MBLAC2-depleted cells. Read fractions are shown for all rDNA-mapping reads and for reads exceeding 250 bp or 1 kb in length. (C) Differential start/end matrix of Pol I-associated rRNA reads comparing siMBLAC2 with siCTR cells. Each bin represents a combination of read start and end positions across the rDNA transcription unit. Red bins indicate read architectures enriched upon MBLAC2 depletion, whereas blue bins indicate those enriched in control cells. Annotated pre-rRNA processing intermediates and mature rRNA regions are indicated. (D) Quantification of read starts near annotated rRNA processing sites. Mature rRNA start sites were excluded from the analysis to focus on internal processing-associated read starts. Values are expressed as CPM within a ±25-nt window around each annotated site. (E) Long-read coverage across the first 2 kb of the downstream intergenic spacer (IGS). Reads overlapping consecutive 1-kb IGS bins were normalized to total rDNA-plus-IGS mapped reads. Increased IGS signal following MBLAC2 depletion is consistent with abnormal persistence or readthrough of Pol I beyond the 3′ end of the transcription unit. The start/end matrix analysis in (C) was performed using a custom workflow adapted from the pre-rRNA processing-site annotation framework described by Pastore et al. Read start and end coordinates were extracted from rDNA-mapped POINT-nano alignments, binned across the rDNA reference, normalized to counts per million mapped reads.

We first assessed whether POINT-nano recovered comparable populations of rRNA-derived molecules in control and MBLAC2-depleted cells. Because human rDNA repeats are heterogeneous and incompletely represented by any single reference sequence, we focused subsequent analyses on the high-confidence subset of POINT-nano reads mapping to the standard 45S rDNA reference used throughout this study. A larger fraction of total reads mapped to the rDNA reference following MBLAC2 depletion, and this increase was evident among reads exceeding 250 bp and 1 kb (Fig. 5B). Thus, POINT-1-nano provided a suitable single-molecule view of Pol I-associated pre-rRNA molecules.

We next examined read start and end positions across the rDNA transcription unit. In control cells, the resulting read architectures were broadly compatible with known pre-rRNA precursors and processing products. In particular, the prominent population of reads extending from the 5’ terminal 01 region into 18S, together with molecules spanning from the 5.8S region towards 28S, resembles the precursor-enriched nuclear rRNA profile. This contrasts with mature rRNA nuclear or cytoplasmic profiles as reported by long-read fractionation studies. This supports the recovery of Pol I-associated nascent and co-transcriptional processing of pre-rRNA by POINT-1^43^. Notably, MBLAC2 depletion produced a distinct pattern in the differential start/end matrix, with enrichment of read architectures extending across spacer-containing regions and toward the 3′ portion of the 47S precursor (Fig. 5C). The corresponding raw start/end matrix showed that these were not isolated events, instead reflecting a broader shift in the population of Pol I-associated RNA molecules (Supplementary Fig. 5A). These observations are consistent with accumulation of incompletely processed and extended pre-rRNA intermediates following loss of MBLAC2.

A characteristic feature of pre-rRNA maturation is the generation and subsequent processing of RNA ends at defined cleavage sites. We therefore quantified read starts near annotated processing positions. MBLAC2 depletion reduced the abundance of reads initiating at several internal processing positions, including sites 3, E1, E2, C, 2, 4 and 02 (Fig. 5D). These sites are normally associated with successive maturation steps within the pre-rRNA precursor. Their reduced representation following MBLAC2 depletion is consistent with inefficient endonucleolytic processing and/or impaired downstream exonucleolytic turnover of these intermediates. By contrast, analysis including all annotated processing sites showed that read starts at sites 1, B1 and 3′ were retained or enriched upon MBLAC2 depletion (Supplementary Fig. 5B). Thus, the MBLAC2 depletion defect is unlikely to reflect a general loss of pre-rRNA cleavage. Rather, it appears to alter the processing of a subset of internal intermediates, leading to the persistence of RNA species that are normally further matured or cleared.

MBLAC2 depletion analysed by POINT-1 RT-qPCR (Fig. 3C) or POINT 1-seq short-read profiles (Fig. 4E) suggested increased Pol I-associated RNA downstream of the 3′ETS. We therefore determined whether POINT-nano also reveals the existence of extended transcripts within the intergenic spacer. Indeed, long-read coverage increased across the first two kilobases of the IGS following MBLAC2 depletion (Fig. 5E). A related increase was detected by short-read POINT-1 sequencing (Supplementary Fig. 5C). Moreover, quantification of reads overlapping the wider IGS region showed increased IGS-containing RNA in Pol I-associated samples following MBLAC2 depletion. Instead, the corresponding Pol II-associated RNA profile showed a distinct and less pronounced pattern (Supplementary Fig. 5E). Thus, the downstream IGS signal is most consistent with a high-level persistence or extension of Pol I-associated RNA beyond the rRNA TU 3′ end.

Together, these long-read data provide direct molecular evidence that MBLAC2 is required to prevent accumulation of aberrant pre-rRNA intermediates. By resolving individual transcripts, POINT-nano shows that MBLAC2 depletion increases rDNA-mapping reads, enriches RNA ends near processing sites and promotes accumulation of extended molecules reaching downstream IGS regions. These findings all combine to predict a mechanism by which MBLAC2 contributes to co-transcriptional rRNA quality control. This acts to enforce efficient processing and release of nascent Pol I transcripts.

## Discussion

Our results support a model (Fig. 6) in which MBLAC2 acts within a nucleolar RNA quality-control pathway to coordinate co-transcriptional pre-rRNA processing with Pol I transcription regulation. RNA surveillance of defective nuclear transcripts commonly involves coordination between endonucleolytic processing, exonucleolytic turnover and the transcription machinery engaged on the template^44,45^. In this context, MBLAC2 depletion causes accumulation of spacer-containing pre-rRNA intermediates, altered processing-site read profiles and readthrough of Pol I into downstream IGS regions. Together, these observations demonstrate that MBLAC2 is required for efficient maturation of nascent pre-rRNA and for its release from the rDNA transcription unit.

**Figure 6.**
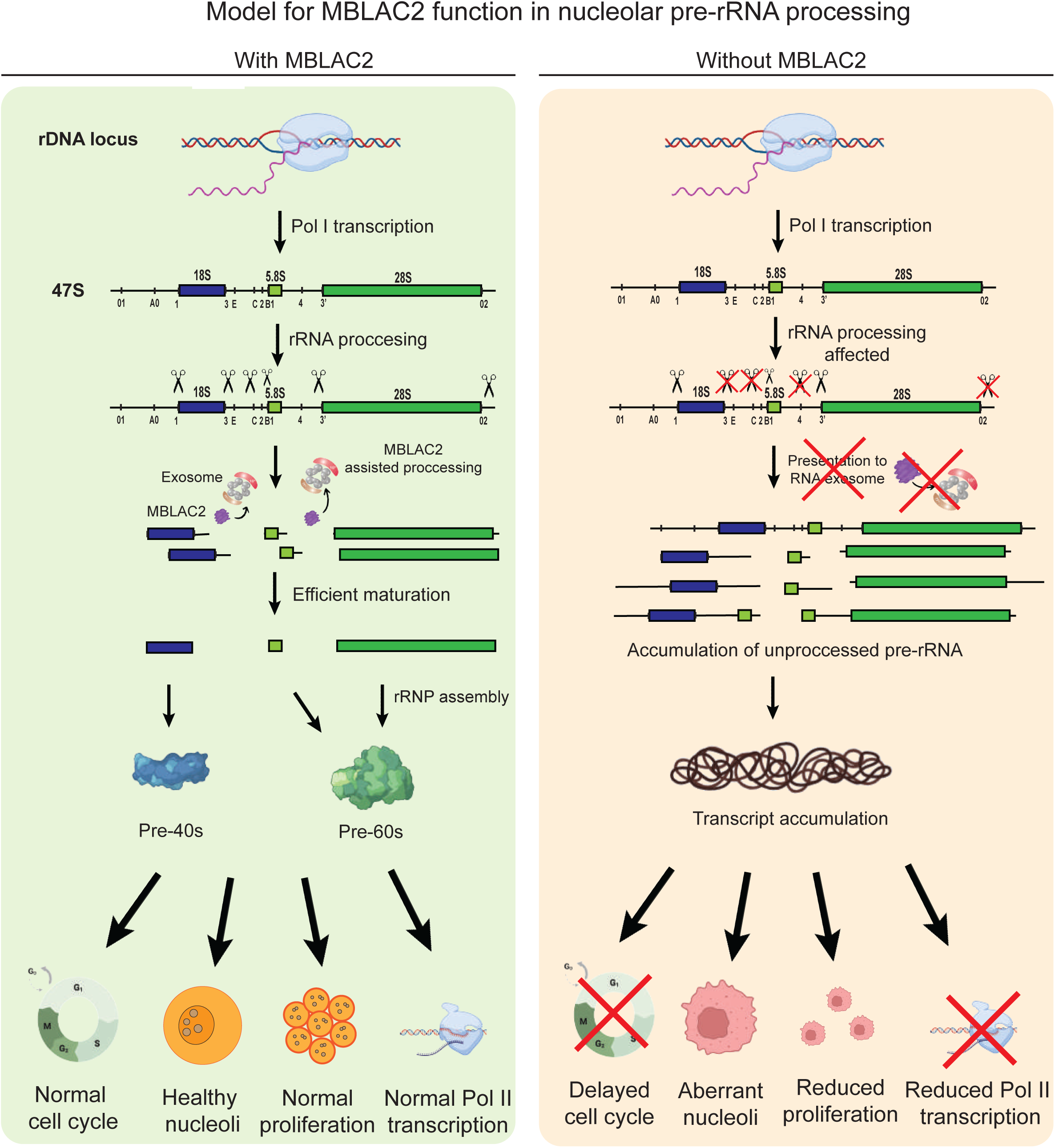
MBLAC2 coordinates co-transcriptional pre-rRNA processing and nucleolar integrity. Working model for MBLAC2 function in nucleolar pre-rRNA processing and transcriptional homeostasis. Under normal conditions, MBLAC2 associates with the Pol I transcriptional environment and promotes efficient co-transcriptional processing of the 47S pre-rRNA. This facilitates removal of transcribed spacer sequences, limits the accumulation of aberrant intermediates and supports productive rRNA maturation and ribosomal subunit assembly. MBLAC2 may also contribute to the generation or presentation of pre-rRNA intermediates for further processing or turnover by the RNA exosome. In the absence of MBLAC2, spacer-containing and extended Pol I-associated transcripts accumulate, including RNA extending into downstream intergenic spacer regions. These defects are accompanied by altered nucleolar architecture, delayed G1/S progression, reduced proliferation and decreased global Pol II-associated nascent transcription. MBLAC2 is therefore proposed to coordinate pre-rRNA processing with Pol I transcript homeostasis, nucleolar organisation and broader cellular transcriptional output.

We propose that under normal conditions, MBLAC2 acts in the Pol I transcriptional environment to promote processing of the 47S precursor and to limit the persistence of aberrant intermediates. By these means MBLAC2 may facilitate the generation of RNA species that undergo further maturation or turnover by the RNA exosome. This would couple early processing events with downstream RNA surveillance and help maintain the balance between rRNA synthesis, processing and nucleolar organisation. Such coupling between rRNA processing, ribosome assembly and Pol I transcription has been described in other systems^46,47^. Upon MBLAC2 depletion, this coordination is disrupted. Spacer-containing pre-rRNA molecules accumulate, internal processing is altered and extended Pol I-associated transcripts persist toward the downstream IGS. These changes are accompanied by the enlargement of fewer nucleoli, altered nucleolar protein abundance, delayed G1/S progression, induction of p21 and reduced global Pol II nascent transcription. Nucleolar disruption and defective ribosome biogenesis are well established triggers of p53/p21-associated cell-cycle control and broader transcriptional adaptation^21,48^. Thus, defective pre-rRNA processing is not confined to the Pol I transcription unit but is associated with a broader change in nucleolar and transcriptional homeostasis^20^. In effect MBLAC2 acts to define a specific nuclear quality control process within the nucleolus. This is dedicated to the efficient and accurate production of mature rRNA ready for assembly into the small and large ribosome subunits.

A central concept emerging from this work is that MBL-domain proteins are organised according to transcript class and processing context. CPSF73 catalyses cleavage during pre-mRNA, INTS11 participates in noncoding RNA processing and transcriptional attenuation, and ELAC2 functions in tRNA maturation^12–14,17^. MBLAC1 has also been linked to histone RNA metabolism, consistent with functional diversification within this protein family^18^. In this context, MBLAC2 fills a distinct nucleolar role. It is functionally separated from MBLAC1 by its lack of detectable association with SLBP, its enrichment in nuclear and chromatin-associated fractions, and the specific nucleolar defects caused by its depletion. These observations argue that MBLAC2 is not a broadly acting MBL-domain nuclease, but rather a factor linked to the processing environment of nascent ribosomal RNA.

The phenotype caused by MBLAC2 depletion is consistent with defective ribosome biogenesis rather than a general consequence of cell-cycle arrest. MBLAC2-depleted cells show enlarged and fewer nucleoli, altered abundance of nucleolar proteins, delayed G1/S progression and reduced cell proliferation. These features resemble, although are generally milder than those observed following depletion of PES1, a ribosome-biogenesis factor required for large ribosomal subunit maturation^31,32^. By contrast, depletion of MBLAC1 or CPSF73 does not reproduce the same nucleolar phenotype. Thus, the consequences of MBLAC2 loss are more consistent with perturbation of nucleolar RNA metabolism than with a secondary response to altered proliferation or general disruption of RNA processing.

At the molecular level, several independent approaches indicate that MBLAC2 depletion alters early pre-rRNA processing. Northern blotting revealed accumulation of spacer-containing pre-rRNA species, while 4sU labelling and POINT-1 RT-qPCR showed increased nascent and Pol I-associated RNA across the 5′ETS, ITS, 3′ETS and downstream IGS regions. These observations indicate that loss of MBLAC2 alters the normal processing and turnover of newly synthesised pre-rRNA. This places the role of MBLAC2 at an early stage of ribosome biogenesis, when processing, folding and assembly of the 47S precursor remain closely coupled.

Our data further indicate that defective processing is associated with altered Pol I transcript homeostasis. MBLAC2 depletion increased Pol I-associated RNA across the transcription unit and in downstream IGS regions, with a relative enrichment of POINT- 1 reads over the 3′ETS. Long-read POINT-nano analysis extended these observations by resolving altered start/end architectures of individual Pol I-associated RNA molecules. MBLAC2-depleted cells accumulated extended and incompletely processed molecules, showed altered representation of several internal processing sites, and displayed increased long-read coverage within the first kilobases of the IGS. Together, these data are consistent with impaired coordination between pre-rRNA processing, transcript release and Pol I termination. Previous studies have shown that rRNA processing, ribosome assembly and Pol I transcription are mechanistically linked^3,37,46^. Our findings extend this principle to human cells by identifying MBLAC2 as a factor required to maintain this coupling.

The POINT-nano data are particularly informative because they show that the phenotype is not limited to increased signal at selected regions of the rDNA repeat. Rather, MBLAC2 depletion changes the distribution of Pol I-associated transcripts across the rRNA TU. The differential behaviour of individual processing-associated positions suggests that MBLAC2 depletion does not produce a uniform change across all annotated rRNA ends. Instead, it alters the distribution of Pol I-associated pre-rRNA intermediates. Loss of MBLAC2 appears to alter processing of a subset of internal intermediates and their subsequent maturation or turnover. This interpretation is consistent with a role for MBLAC2 in generating, stabilising or presenting specific pre-rRNA intermediates for downstream processing.

An additional major finding is that the status of pre-rRNA processing influences global Pol II transcriptional output. Although rRNA is transcribed by Pol I, ribosome biogenesis depends on the coordinated activity of all three nuclear RNA polymerases. Pol II produces mRNAs encoding ribosomal proteins and assembly factors, whereas Pol III produces 5S rRNA and tRNAs required for ribosome function^1,2,5^. MBLAC2 depletion reduced global Pol II-associated nascent RNA across protein-coding genes, and a related effect was observed following PES1 depletion. Importantly, this reduction was not accompanied by broad redistribution of Pol II-associated reads across genomic compartments. Thus, defective pre-rRNA processing appears to reduce the overall amplitude of Pol II transcription rather than extensively altering the genomic distribution of Pol II.

This distinction may help explain why acute Pol I inhibition and chronic defects in pre-rRNA processing produce different transcriptional states. Acute treatment with CX- 5461 reduced Pol I-associated RNA across the rDNA unit, whereas MBLAC2 or PES1 depletion led to persistence of abnormal Pol I-associated transcripts, nucleolar reorganisation and transcriptional adaptation. These conditions are therefore not equivalent. The former reflects reduced production of nascent rRNA while the latter causes failure to efficiently process and release newly synthesised pre-rRNA.

Consistent with this functional interdependence, short-term inhibition of Pol I, Pol II or CDK9-dependent Pol II elongation disrupted the normal distribution of UBF and Nucleolin. These observations indicate that nucleolar organisation is sensitive not only to Pol I activity, but also to the wider transcriptional state of the cell.

RNA-seq analysis further provided a link between MBLAC2-dependent pre-rRNA processing and RNA surveillance. MBLAC2 depletion induced a transcriptional programme associated with cell-cycle delay and ribosome-biogenesis stress, including induction of CDKN1A/p21. At the same time, levels of EXOSC3, encoding a core component of the nuclear RNA exosome, were reduced. Native co-immunoprecipitation showed that MBLAC2 associates with RNAPOL1A and EXOSC3, placing it in proximity to both the Pol I transcriptional environment and a major RNA-surveillance machinery. The nuclear exosome is required for the maturation and turnover of pre-rRNA intermediates and other unstable nuclear transcripts^45,49^. MBLAC2 may therefore act upstream or be coordinated with the exosome machinery. One possibility is that MBLAC2 promotes formation of RNA intermediates that can subsequently undergo exonucleolytic maturation or decay. Alternatively, MBLAC2 may contribute to assembly of a processing environment that acts to couple cleavage, transcript release and surveillance of aberrant precursors.

Recent work has implicated MBLAC2 in the suppression of rDNA transcription following nucleolar DNA damage^36^. Our findings now raise the possibility that its basal role in pre-rRNA processing and RNA surveillance may also influence how Pol I transcription is regulated under damage-associated stress. More broadly, the identification of this layer of nucleolar RNA quality control raises questions related to transcription-coupled repair at Pol I-transcribed genes. Here the relationship between transcriptional arrest, RNA processing and rDNA repair remains unclear^50^.

Together, our findings identify MBLAC2 as a nucleolar MBL-domain factor required for co-transcriptional pre-rRNA processing and Pol I-associated RNA regulation. More broadly, they support a model in which MBL-domain proteins form a compartmentalised RNA quality-control network in which distinct factors safeguard different classes of abundant or unstable transcripts. Within this family, MBLAC2 links the processing state of the most abundant nuclear transcript, rRNA, to nucleolar organisation, cell-cycle control and global Pol II transcriptional output.

### Limitations

A limitation of the present study is that direct biochemical cleavage assays using physiologically relevant human pre-rRNA substrates remain technically challenging. Human Pol I transcripts are abundant, heterogeneous in sequence, GC-rich, highly structured and rapidly assembled into ribonucleoprotein complexes. In addition, rDNA repeats are heterogeneous and are processed within the specialised environment of the nucleolus. We therefore cannot distinguish between MBLAC2 directly cleaving a defined pre-rRNA substrate, facilitating the activity of another nuclease, or promoting access of processing intermediates to the exosome. Nevertheless, our combined genetic, biochemical and transcriptomic evidence supports a role for MBLAC2 in the processing of nascent pre-rRNA.

## Acknowledgments

We thank A. Kornblihtt and N.J.P. group members for critical discussions. We are also grateful to the Don Mason Flow Cytometry Facility at the Dunn School, as well as R. Hedley and Vasiliki Tsioligka for assistance. We would like to thank Alan Wainman and the Dunn School Bioimaging Facility for expert advice and access to microscopes.

L.E.M. has received ongoing support from Linacre College, through a Paul Nurse Junior Research Fellowship. C.J.S thanks Cancer Research UK for support (CRUK/ A24759). This project was supported by funding from a Wellcome Trust Investigator Award (107928/Z/15/Z) to N.J.P.

## Declaration of interest

L.E.M., C.J.S. and N.J.P. declare that they have no competing financial or non-financial interests related to the work described in this study. None of the authors has a commercial, advisory, patent-related or other relationship that could be perceived as influencing the interpretation or presentation of these findings.

## Author contributions

L.E.M. performed most of the experiments and analyses. AS.M. performed additional experiments. R.Sm. performed the microscopy experiments and image analyses. R.S.L. performed initial processing and filtering of sequencing datasets and provided intellectual input. C.J.S provided invaluable expertise in the human MBL superfamily. N.J.P. supervised the work. L.E.M. and N.J.P. wrote the manuscript with input from all authors.

## Lead contact

Further information and requests for resources and reagents should be directed to and will be fulfilled by the lead contact, Dr. Luciano E. Marasco

## Materials availability

All unique/stable reagents generated in this study are available from the lead contact upon request.

## STAR Methods

### Cell culture and treatments

HCT116 and HeLa cells were grown in Dulbecco’s modified Eagle’s medium (DMEM) containing 4.5 g of glucose and 10% fetal bovine serum (Gibco) at 37 °C. Cells were plated at a density of 2.10^5 cells per well in 6-well plates 24 hr before transfection. siRNA (25 nM) or plasmid (500 ng) transfections were performed 24 hr after cells were plated, using 3 μl of Lipofectamine 2000 (Thermo Fisher Scientific) per well in 6-well plates. After 48 hr, RNA Pol I transcription was inhibited with CX-5461 (Merck, 5092650001) at 100 nM for 1 hr. RNA Pol II was degraded with α-amanitin (Merck, A2263) at 1 µg/mL for 4 hr or elongation blocked with DRB (Merck, D1916) at 100 μM for 4 hr. Vehicle-treated cells were used as controls and harvested for downstream procedures. The cell passage number never exceeded ten passages. All harvestings and experiments occurred at 70-85% confluency

### RNAi knockdown

Downregulation of MBLAC2, MBLAC1, CPSF73, PES1, HAGH, ETHE1 or control siRNA was performed using ON-TARGET plus SMARTpool siRNA oligonucleotides (Dharmacon). siRNA oligos were delivered to cells following the manufacturer’s instructions and allowed to act for 48 hr. Accell siRNA anti-human, non-targeting siRNA (Dharmacon, NC1567415) was used as a control. Knockdown efficiency was assessed by RT-qPCR and/or western blotting where antibodies were available. siRNA sequences are listed in Supplementary Table 1

### Plasmid transfection and overexpression

Plasmid transfections were performed as described above. ORFs encoding MBLAC1 and MBLAC2 were obtained from Origene (RC206960 and RC207867, respectively) and subcloned into the pCI vector. To facilitate immunoprecipitation, MBLAC1 was tagged with FLAG and MBLAC2 with HA. Cells were transfected with empty vector, epitope- tagged MBLAC1, wild-type MBLAC2 or a catalytically impaired MBLAC2 mutant. Cells were harvested 8-16 hr after transfection for immunofluorescence, biochemical fractionation, proliferation assays, Annexin V/PI analysis or co-immunoprecipitation.

### Cell-cycle synchronization and flow cytometry

For cell-cycle analysis, HCT116 cells were synchronized using a double thymidine block.^49^ Cells were treated with 2mM of thymidine for 12 hr, released by washing with fresh medium, treated again with thymidine for 12 hr, and released for 0, 3, 6, 9 and 12 hr before collection. Cells were fixed in 70% ethanol, stained with propidium iodide (Merck, P4170) in the presence of RNase A (Thermo, EN0531), and analyzed by flow cytometry (BD Fortessa X20). Cell-cycle distribution was calculated using FlowJo based on DNA content.

### Annexin V and propidium iodide apoptosis assay

Cells were collected 48 hr after siRNA transfection or 12 hr after plasmid overexpression, washed with PBS and stained with Annexin V - Alexa Fluor 488 conjugate (Thermo Fisher, A13201) and propidium iodide according to the manufacturer’s protocol. Samples were analyzed by flow cytometry. Live, early apoptotic, late apoptotic and dead populations were defined according to Annexin V and PI staining^52^.

### Crystal violet proliferation assay

Cells were seeded at equal density before siRNA transfection or plasmid overexpression and collected at the indicated time points. Cells were fixed with methanol, stained with crystal violet, washed and imaged. For quantification, data was analysed and quantified using image J colony area plugin with manual plate input pf K2 =0.15 and K3=0.15. After bound dye was solubilized in SDS solution and absorbance was measured at 562 nm. Values were normalized to control where indicated.

### Immunofluorescence microscopy

For Figure 2 and Supplementary Figure 2 and 4, siRNA mediated knockdown in HCT116 cells was performed using Lipofectamine RNAiMAX (Invitrogen) according to the manufacturers’ instructions in a 24-well imaging plates (Miltenyi Biotec) 72 hr prior to fixation. Media was removed from the cells, and they were washed once with PBS. PBS was removed and cells were fixed in 4% PFA in PBS for 15 min at room temperature. Cells were washed twice with PBS after fixation and then permeabilised with 0.2% Triton-X in PBS for 5 min with gentle rocking. Cells were washed once more with PBS and incubated in blocking buffer (3% BSA,w/v, in PBS + 0.2% Tween 20) for 60 min. Cells were incubated in primary antibody (Nucleolin, ab22758, 1:1500; UBF, sc- 13125,1:500; FLAG, F7425, 1:200; MBLAC2, HPA060264, 1:100) diluted in blocking buffer overnight at 4°C. Primary antibody was removed from the wells and the cells were washed three times with PBS + 0.1% Triton. Secondary antibody was diluted 1:500 in blocking buffer with 1.25 µg/mL Hoechst 33342 (Thermo Fisher Scientific) for 1 hr at room temperature with gentle rocking. Cells were washed three times with PBS + 0.1% Triton prior to imaging. Figure images were captured on an Olympos IX-83 inverted microscope equipped with a Yokogawa SoRa spinning disk, with a 60x/1.5 numerical aperture oil-immersion objective, and a Prime 95B scientific complementary metal- oxide semiconductor camera. The fluorophores were excited with lasers at 405 nm and 488 nm, and fluorescence was detected with band-pass filters adapted to the fluorophore emission spectra. Figure (both main and Supplementary) images were obtained using a Zeiss LSM880 inverted confocal microscope equipped with a Plan- APO Chromat 63X /1.4 N.A. oil-immersion objective lens. Fluorophores were excited with 405 nm and 633 nm solid state lasers or a 488 nm argon laser. Fluorescence was detected using GaAsP detectors with collection windows set according to the fluorophore.

Image quantification was performed using CellProfiler using a custom pipeline^53^. Nuclei were identified through Hoescht staining while Nucleoli were identified using Nucleolin.

### Biochemical fractionation

Cells were fractionated using Abcam’s Standard Cell Fractionation Kit (Abcam) Fractions were analysed by SDS-PAGE and western blotting. MBLAC2 distribution was assessed using antibodies against endogenous MBLAC2 or epitope-tagged MBLAC2.

### RNA extraction and RT-qPCR

Total or chromatin RNA was extracted using TRIzol reagent and treated with DNase I to remove any genomic DNA contamination. cDNA synthesis was performed using random hexamers and SuperScript III (Thermo Fisher, 18080085) according to the manufacturer’s protocol. qPCR was carried out using SYBR Green or probe-based chemistry on a Qiagen Rotor-Gene as previously described^54^. Primer sequences are listed in Supplementary Table 2.

### Western blotting

Cells were lysed in 1X protein sample buffer (50mM Tris-HCl pH6.8, 2% SDS, 10% glycerol, 5% β-mercaptoethanol, 0.0025% Bromophenol blue). Protein concentration was determined using BCA (Life Technologies, 23227). Equal amounts of protein were resolved by 4-12% SDS-PAGE (NuPage, Life Technologies) and electroblotted onto nitrocellulose membranes (Merck, GE10600002). Membranes were blocked with milk, incubated with primary antibodies listed in Supplementary Table 1, followed by fluorescent-conjugated secondary antibodies (LI-COR Biosciences, 926-32211 or 926- 32212). Detection was performed using Odyssey instrument (LI-COR Biosciences). Vinculin, tubulin or histone H3 were used as loading controls.

### Native co-immunoprecipitation

Cells expressing tagged MBLAC1 or MBLAC2 were lysed under native conditions in IP/lysis buffer containing 50 mM Tris–HCl (pH 7.6), 150 mM NaCl, 1% NP-40, 1 mM EDTA, 1 mM EGTA, 2% glycerol, protease and phosphatase inhibitor cocktails and 50 U/mL Benzonase. Lysates were cleared by centrifugation and incubated with anti- HA/anti-FLAG antibody-coupled beads for 4 hr at 4°C. Input, flow-through and immunoprecipitated fractions were analyzed by western blotting. Co- immunoprecipitation was used to assess association of MBLAC1 or MBLAC2 with SLBP, RNAPOL1A and EXOSC3. As a specificity control for POINT-1, native RNAPOL1A immunoprecipitation was tested for co-inmunoprecipitation of RNAPOL2 and RNAPOL3A.

### Northern blotting

Total RNA was extracted from control and MBLAC2-depleted cells, 5-10 ug were resolved on formaldehyde-agarose gels, then transferred to Amersham Hybond-N+ membranes (Cytiva, RPN203B). Biotinylated probes targeting rRNA regions (Supplementary Table 2) were used for hybridization. Membranes were blocked and then incubated with IRDye 800CW Streptavidin (LI-COR, #926-32230) prior to detection. Signals were imaged using an Odyssey instrument (LI-COR). Methylene blue staining and/or U6 snRNA served as loading controls. The full protocol is available upon request.

### Puromycin (SUNSET) assay

Global translation was monitored by puromycin incorporation as described^55^. Cells were incubated with puromycin at 10 uM for 10 minutes before harvest. Protein extracts were analyzed by western blotting using anti-puromycin antibody. Vinculin and tubulin were used as loading controls.

### 4sU labelling and biotinylation

4sU labelling and biotinylation were performed as described^56^. For HCT116 cells, two 100 mm dishes were labelled with 4sU for 5 min, followed by RNA extraction, biotinylation and streptavidin purification as described.

### POINT-seq and POINT-nano

POINT-seq and POINT-nano was performed as described.^38^ For POINT-1, DNase I digestion time was doubled compared to the standard protocol and the POLR1A (D6S6S) rabbit monoclonal antibody (Cell Signaling, #24799) was used for immunoprecipitation. Libraries were prepared using the NEBNext Ultra II Directional RNA Library Prep Kit for Illumina, deep sequencing was conducted on a NovaSeq 6000 platform (Illumina) by Novogene UK. POINT-Nano libraries were prepared using the SQK-RNA004 kit (ONT) following the manufacturer’s protocol and then sequenced on MinION using MinION RNA Flow Cell (ONT, MIN004RA).

### rDNA and IGS reads analysis

For short-read POINT-seq and POINT-nano analyses, reads were mapped to the human rDNA locus 45SN1 (equivalent to 47S; Gene ID: 106631777; NW_021160023.1:480347-493697), which contains the 47S transcription unit and downstream intergenic spacer (IGS).

### Data processing

Data processing PolyA-selected RNA-seq, POINT-seq and POINT-nano datasets were analyzed using bioinformatic pipelines adapted from those described by Marasco et al^54^. Briefly, sequencing reads were subjected to quality control and adapter trimming before alignment to either the human reference genome or the rDNA reference described above, depending on the analysis. PolyA-selected RNA-seq reads were aligned to the human genome and used for gene-level quantification, differential expression analysis and gene ontology enrichment. Pol II POINT-seq datasets were used to generate metagene profiles and genomic feature distributions, whereas Pol I POINT-seq datasets were mapped to the rDNA/IGS reference to generate rDNA coverage profiles and quantify IGS readthrough. For rDNA-specific analyses, short- read POINT-seq and POINT-nano reads were mapped to the human rDNA locus 45SN1, equivalent to the 47S rRNA transcription unit, together with its downstream intergenic spacer. For POINT-nano datasets, rRNA read start and end positions were extracted from long-read alignments and used to generate start/end matrices. Enrichment of read starts near annotated rRNA processing sites was calculated using scripts described in Pastore et al^43^.

## Supplementary Figure Legends

**Supplementary Figure 1.**
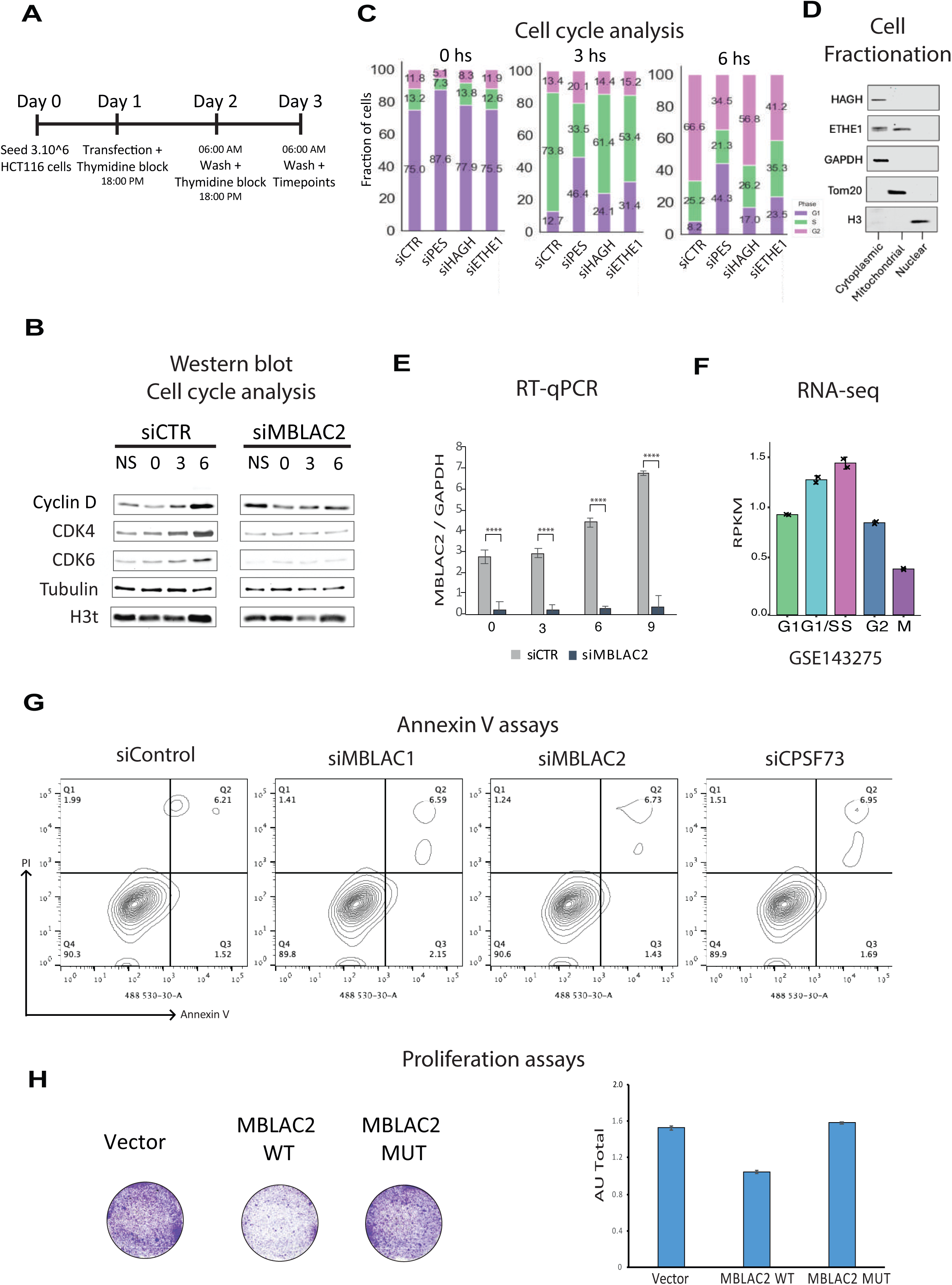
MBLAC2 depletion perturbs cell-cycle progression without inducing an apoptotic response. (A) Experimental design for cell-cycle synchronization. HCT116 cells were transfected with the indicated siRNAs, synchronized by double-thymidine block and released for the indicated times before collection. (B) Western blot analysis of G1/S regulators in control and MBLAC2-depleted cells collected before synchronization (NS) or at 0, 3 and 6 h after release. Cyclin D, CDK4 and CDK6 were analysed, with tubulin and total histone H3 (H3t) shown as loading controls. (C) Cell-cycle distribution following depletion of selected MBL-domain proteins or PES1. DNA content was measured by propidium iodide staining and flow cytometry. The percentages of cells in G1, S and G2/M phases are shown for each condition and time point. (D) Subcellular fractionation of HCT116 cells into cytoplasmic, mitochondrial and nuclear fractions, followed by immunoblotting for HAGH and ETHE1. GAPDH, TOM20 and histone H3 were used as cytoplasmic, mitochondrial and nuclear fraction markers respectively. (E) RT–qPCR analysis of MBLAC2 depletion during the cell-synchronization experiment. MBLAC2 mRNA levels were normalized to GAPDH and expressed relative to siCTR. (F) Re-analysis of a published RNA-seq dataset showing MBLAC2 transcript abundance across the indicated cell-cycle phases. RPKM values are shown. Dataset accession: GSE143275. (G) Annexin V/propidium iodide flow-cytometry analysis following depletion of MBLAC1, MBLAC2 or CPSF73. Representative contour plots are shown. (H) Crystal-violet proliferation assay following expression of empty vector, wild-type MBLAC2 or the catalytically impaired MBLAC2 mutant. Representative wells are shown on the left, and quantification of solubilized crystal-violet staining is shown on the right. Quantitative data are presented as mean ± s.e.m. from three independent biological replicates. Cell-cycle data in (C) and RT–qPCR data in (E) were analysed by two-way ANOVA followed by Šídák’s multiple-comparisons test. Proliferation data in (H) were analysed by one-way ANOVA followed by Tukey’s multiple-comparisons test. The RNA-seq data in (F) are presented descriptively. Western blots and Annexin V/propidium iodide plots are representative of independent experiments. *P < 0.05; *P < 0.01; *P < 0.001; *P < 0.0001; ns, not significant.

**Supplementary Figure 2.**
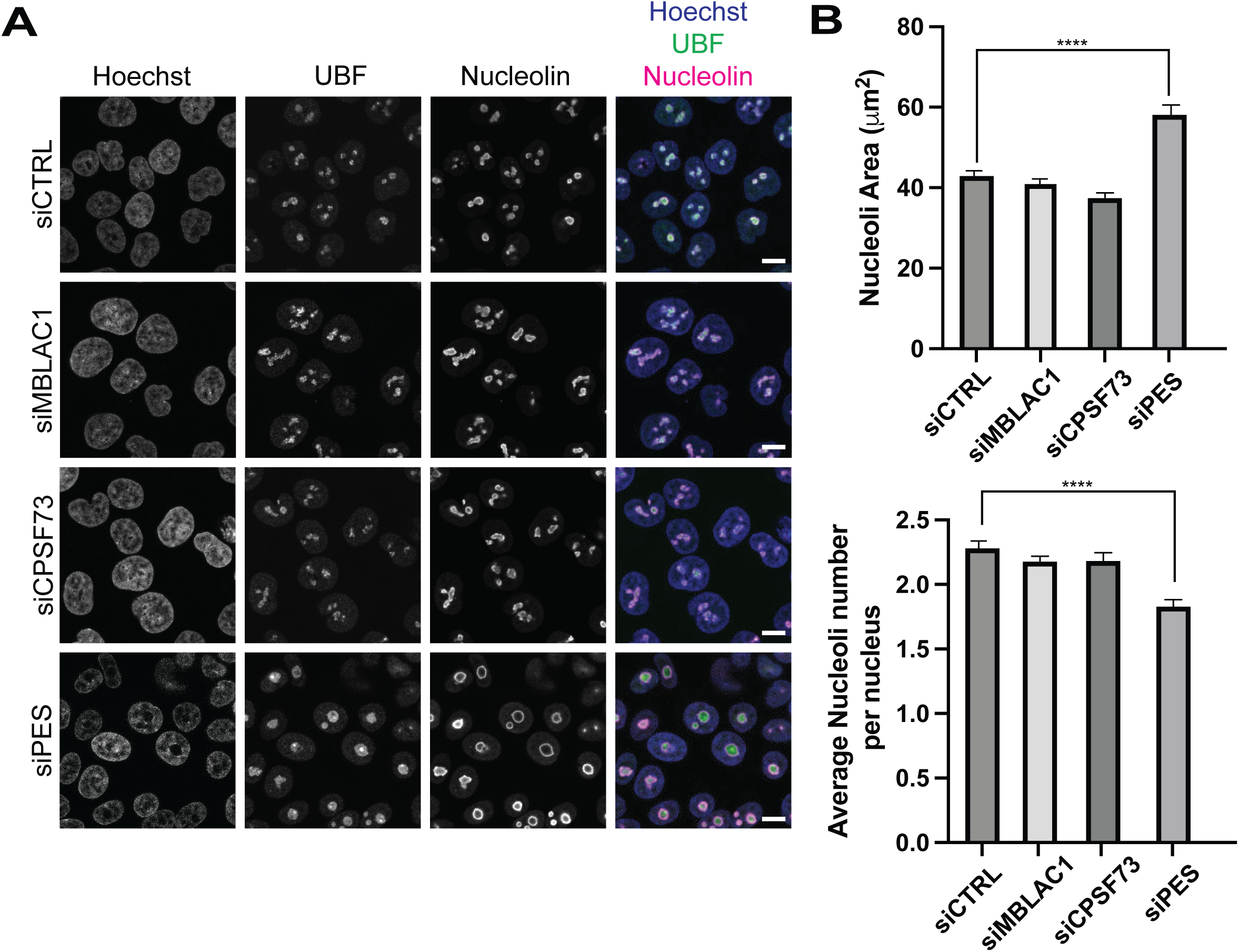
MBLAC1 depletion does not phenocopy the nucleolar defects caused by MBLAC2 depletion. (A) Immunofluorescence analysis of nucleolar morphology following depletion of MBLAC1, CPSF73 or PES. Cells were stained for UBF and nucleolin, and nuclei were counterstained with Hoechst. PES1 depletion is shown as a positive control for altered nucleolar morphology. (B) Quantification of nucleolar area (top) and average nucleolar number per nucleus (bottom) from the experiment shown in (A). Quantification is shown for one representative experiment from three independent biological replicates, with 209–286 cells analysed per condition. Data were analysed by one-way ANOVA followed by Dunnett’s multiple-comparisons test against siCTR. Bars represent mean ± s.e.m. Images are representative of three independent experiments. ****P < 0.0001.

**Supplementary Figure 3.**
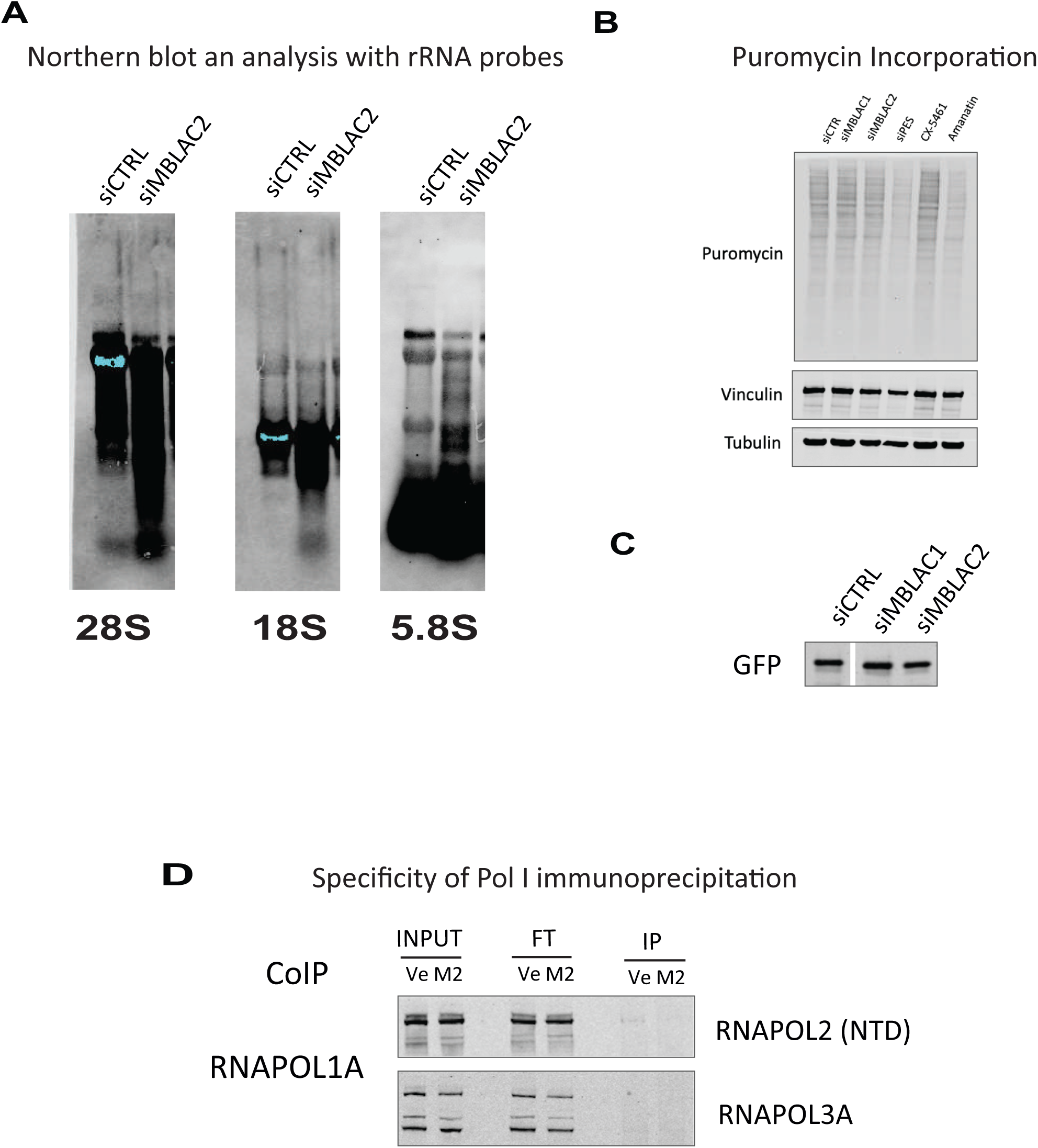
MBLAC2 depletion alters pre-rRNA processing without impairing global translation. (A) Northern-blot analysis of mature rRNA species in control and MBLAC2-depleted cells using probes against 28S, 18S and 5.8S rRNAs. (B) SUNSET assay assessing global translation after the indicated knockdowns or treatments. Puromycin incorporation was detected by western blotting. Vinculin and tubulin are shown as loading controls. (C) Western-blot analysis of GFP expression following depletion of MBLAC1 or MBLAC2, as an additional control for global protein-expression capacity. (D) Specificity control for native RNA polymerase I immunoprecipitation used for POINT-1 analysis. Input, flow-through and immunoprecipitated fractions were analysed for RNA polymerase I and the indicated RNA polymerase II/III markers. Western blots and northern blots are representative of n = 2 independent experiments.

**Supplementary Figure 4.**
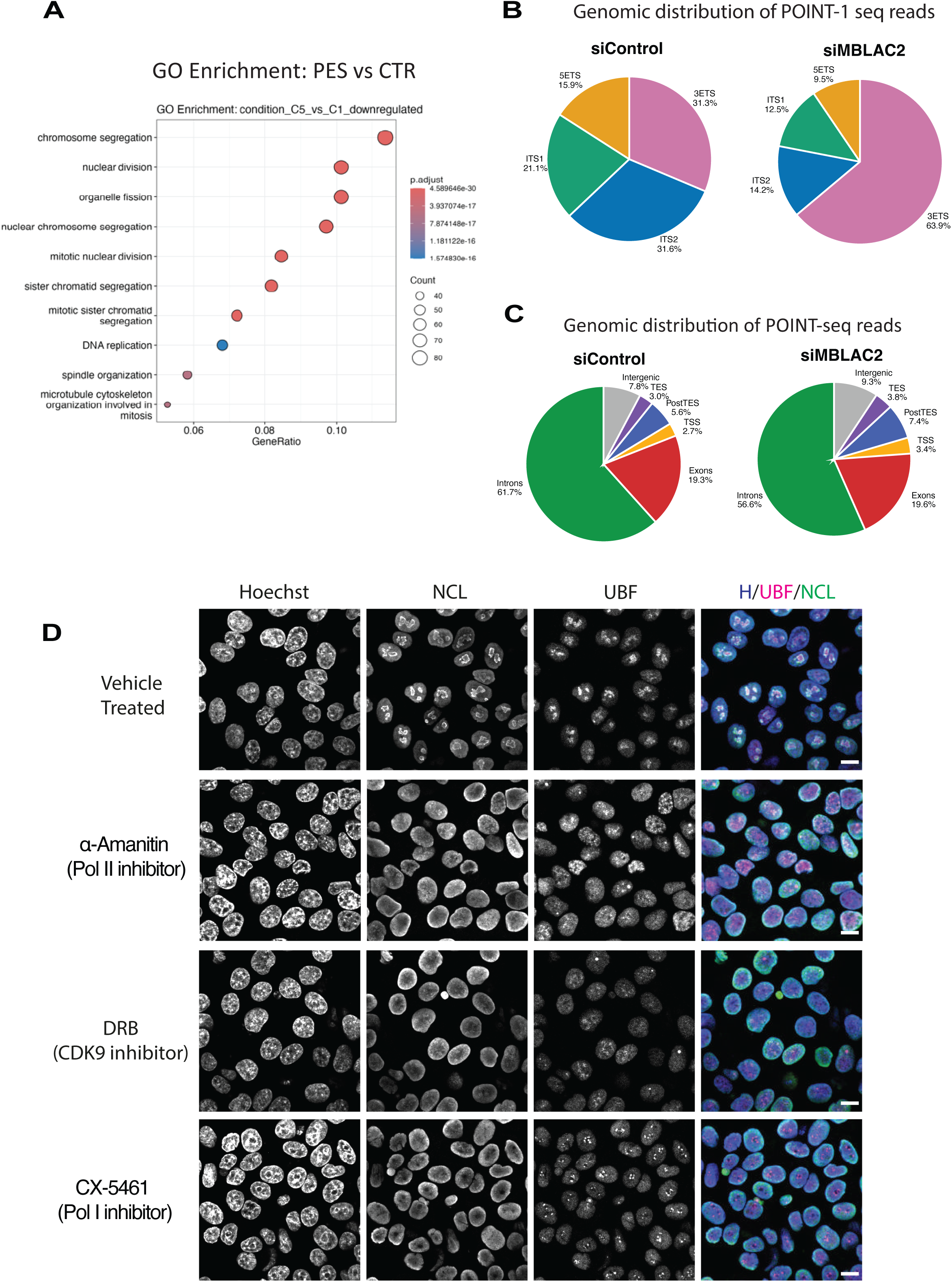
MBLAC2 and PES1 depletion induce related transcriptional and nucleolar-stress responses. (A) Gene ontology enrichment analysis of differentially expressed genes following PES1 depletion relative to siCTR cells. Dot size indicates the number of genes assigned to each term, and colour indicates the adjusted P value. (B) Distribution of POINT-1 seq reads across major regions of the 47S pre-rRNA transcription unit in siCTR- and siMBLAC2-treated cells. Reads were assigned to the 5′ETS, ITS1, ITS2 and 3′ETS regions. MBLAC2 depletion increases the relative contribution of Pol I-associated reads mapping to the 3′ETS region. (C) Distribution of POINT-2 seq reads across genomic features in siCTR and siMBLAC2 treated cells. Reads were assigned to introns, exons, transcription start site (TSS)- proximal regions, transcription end site (TES) regions, post-TES regions and intergenic sequences. (D) Immunofluorescence analysis of nucleolar architecture following short-term inhibition of Pol II with α-amanitin, inhibition of CDK9-dependent transcriptional elongation with DRB or inhibition of Pol I with CX-5461. Cells were stained for UBF and Nucleolin, and nuclei were counterstained with Hoechst. Vehicle-treated cells are shown as controls. Differential-expression analysis in (A) was performed using DESeq2, with P values adjusted using the Benjamini–Hochberg procedure. Gene ontology enrichment analysis was performed using clusterProfiler with false-discovery-rate correction. Read distributions in (B) and (C) are shown as normalized proportions and are presented descriptively. Images in (D) are representative of n = 2 independent experiments.

**Supplementary Figure 5.**
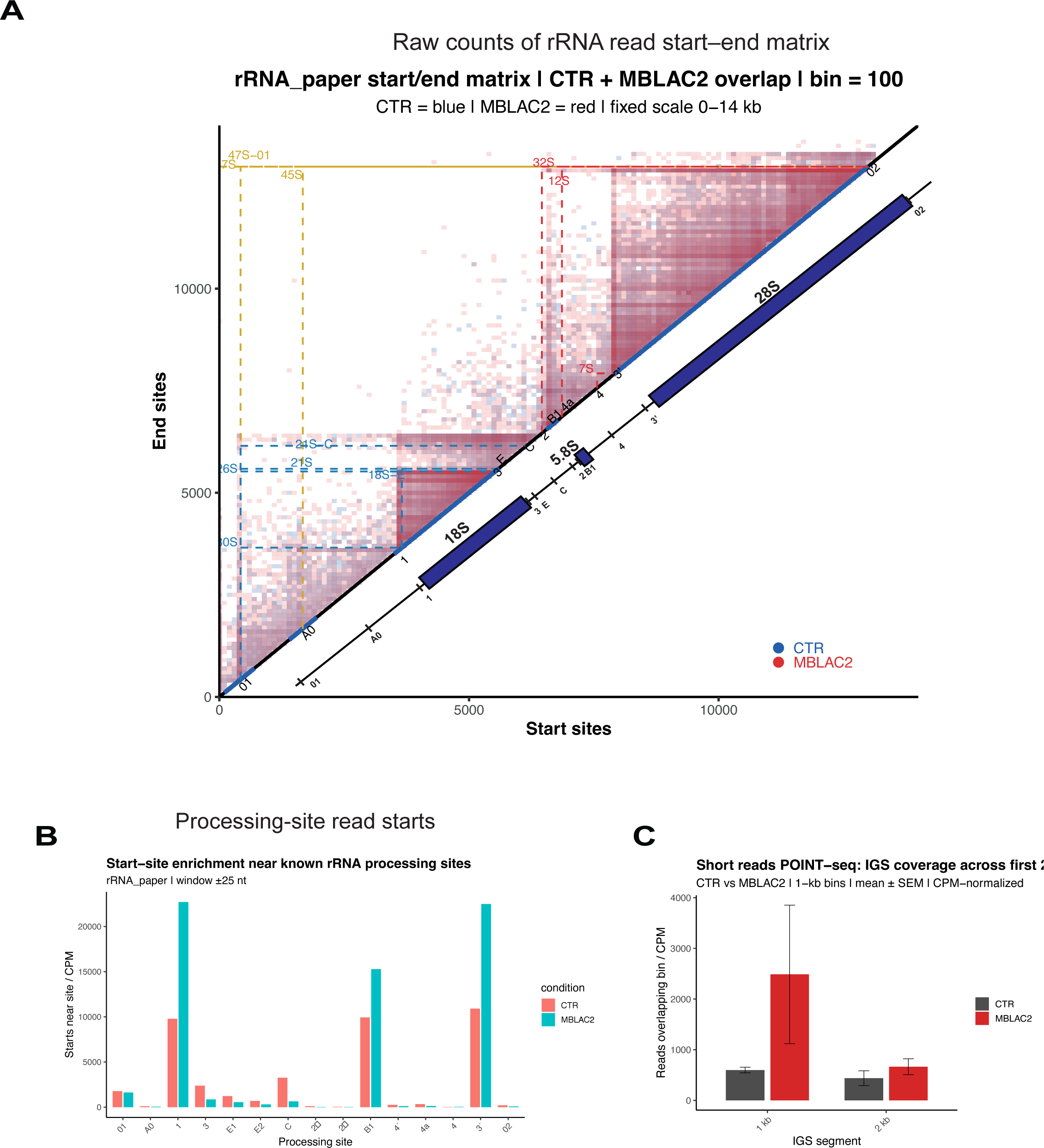
Additional long-read analyses support aberrant processing and IGS-associated readthrough after MBLAC2 depletion. (A) Raw start/end matrix of rDNA-mapping POINT-nano reads in control and MBLAC2-depleted cells. Each bin represents the number of long reads defined by their mapped 5′ start and 3′ end coordinates across the rDNA reference. Control and MBLAC2-depleted reads are shown in blue and red, respectively. Annotated pre- rRNA intermediates and mature rRNA regions are indicated. (B) Quantification of POINT-nano read starts near annotated rRNA processing sites. Values are expressed as counts per million within a ±25-nt window around each processing site. This analysis includes: 1, B1 and 3’ cut sites. (C) Short-read POINT-1 coverage across the first two 1-kb bins of the downstream intergenic spacer (IGS) in control and MBLAC2-depleted cells. Panels (A) and (B) are derived from POINT-nano long-read sequencing. Panel (C) is derived from short-read POINT-seq datasets. Where replicate-level data were available, values are shown as mean ± SEM from n = 2 independent biological replicates.

## Notes

### Competing Interest Statement

The authors have declared no competing interest.

